# A cross-disease human microglial framework identifies disease-enriched subsets and tool compounds for microglial polarization

**DOI:** 10.1101/2022.06.04.494709

**Authors:** John F. Tuddenham, Mariko Taga, Verena Haage, Tina Roostaei, Charles White, Annie Lee, Masashi Fujita, Anthony Khairallah, Gilad Green, Bradley Hyman, Matthew Frosch, Sarah Hopp, Thomas G. Beach, John Corboy, Naomi Habib, Hans-Ulrich Klein, Rajesh Kumar Soni, Andrew F. Teich, Richard A. Hickman, Roy N. Alcalay, Neil Shneider, Julie Schneider, Peter A. Sims, David A. Bennett, Marta Olah, Vilas Menon, Philip L. De Jager

## Abstract

Human microglia play a pivotal role in neurological diseases, but few targeted therapies that directly modulate microglial state or function exist due to an incomplete understanding of microglial heterogeneity. We use single-cell RNA sequencing to profile live human microglia from autopsies or surgical resections across diverse neurological diseases and central nervous system regions. We observe a central divide between oxidative and heterocyclic metabolism and identify subsets associated with antigen presentation, motility, and proliferation. Specific subsets are enriched in susceptibility genes for neurodegenerative diseases or the disease-associated microglial signature. We validate subtypes *in situ* with an RNAscope-immunofluorescence pipeline and leverage our dataset as a classification resource, finding that iPSC model systems recapitulate substantial *in vivo* heterogeneity. Finally, we identify and validate candidates for chemically inducing subtype-specific states *in vitro*, showing that Camptothecin downregulates the transcriptional signature of disease-enriched subsets and upregulates a signature previously shown to be depleted in Alzheimer’s.

## Introduction

Microglia are the resident parenchymal myeloid population of the central nervous system (CNS)^1^. They are well-known for their plasticity. Optimized as sensors and first responders in their environment, they can rapidly disengage from key homeostatic functions to fulfill different specialized roles, such as antigen presentation, pathogen response, and sculpting their cellular milieu^2, 3^. They have pivotal roles in both CNS development^2^ and diseases ranging from Alzheimer’s disease (AD)^4^ to multiple sclerosis (MS)^5^. We are only beginning to understand their spatial, temporal, and functional complexity, with much of the current work performed in mice^6, 7^. However, a pivotal question remains poorly explored: what is the full extent of microglial heterogeneity across different diseases, regions, and the life course in humans? In the past decade, many groups have demonstrated shifts in the human microglial transcriptome with age^8, 9^ or in association with AD pathology^10, 11^. However, structured evaluations of human microglial heterogeneity at the single cell level have only recently been applied to a limited set of contexts^4, 11–18^. Further, most of these studies analyze only a modest number of samples or use single-nucleus profiling, which, as it does not capture cytoplasm, may capture slightly different genes than single-cell, especially in microglia^19, 20^. As a result, our understanding of the range of states that live human microglia can attain as well as their trajectories of state transition remains limited. Analyzing data captured in different contexts in a single framework is essential to interpret results across diseases and studies.

In this study, we aimed to (1) generate a broad reference of microglial transcriptional profiles across neurodegenerative diseases that would capture as much of the diversity of microglial states as possible and (2) illustrate the utility of this resource to annotate model systems and identify tool compounds for modulating human microglial states. Using cold, enzyme-free, mechanical dissociation, we purified live CD45^+^ cells and collected single-cell RNA sequencing (scRNA-seq) data from a diverse set of CNS regions and clinicopathologic states affecting both men and women. We identified 12 microglial subpopulations represented across all diseases and regions; importantly, our study was not designed to characterize microglia in a particular disease, but rather to sample as many different conditions as possible. We propose trajectories of cell state transitions between microglial subsets in our dataset, identifying a central metabolic shift between oxidative and heterocyclic metabolism, identified disease-gene-enriched microglial subsets, microglial subsets that express high levels of the DAM transcriptional program^6^, and subsets associated with immune activation. Using our new subtype signatures, we developed a joint protein-RNA staining protocol to identify microglial subsets *in situ* and demonstrate morphological shifts associated with expression of different hallmark genes. We then used this resource to classify microglia profiled in previous studies and evaluate the degree of microglial diversity found in induced pluripotent stem cell (iPSC)-derived microglial model systems. Finally, we leveraged the Connectivity Map (CMAP)^21, 22^ to identify chemical perturbations predicted to drive subtype-specific signatures and cell state transitions, and we validated these predictions *in vitro* at the RNA and protein levels. Ultimately, we provide a resource that explores human microglial heterogeneity across regions and diseases and a series of tools for classifying, evaluating, and manipulating microglial model systems, bringing us closer to the goal of microglial modulation in humans.

## Results

### Overview of our samples and analytical approach

Our sample collection encompassed fresh autopsy samples from individuals with both early and late-onset AD, mild cognitive impairment (MCI), amyotrophic lateral sclerosis (ALS), frontotemporal dementia (FTD), Parkinson’s disease (PD), progressive supranuclear palsy (PSP), diffuse Lewy body disease (DLBD), MS, Huntington’s disease (HD), and stroke, as well as samples from an individual without a diagnosis of neurologic disease **(Graphical Abstract)**. Our cohort also includes surgical resections from temporal lobe epilepsy (TLE) and a dysembryoplastic neuroepithelial tumor (DNET). These samples were derived from a wide array of brain regions: anterior watershed white matter, frontal cortex (BA9/46), primary motor cortex (BA4), temporal cortex (BA20/21), occipital cortex (BA17/18/19), hippocampus, thalamus, substantia nigra, facial motor nucleus, and spinal cord. As our workflow and budget limited the number of regions that could be processed in parallel for each brain, we chose BA9 as a reference in most cases and sampled other regions where possible. As individuals without any diagnosed pathology (“control” subjects) rarely come to autopsy, only one was sampled. Our workflow included the use of a previously reported cold, enzyme-free mechanical dissociation approach for isolation of live human microglia and leukocytes^4, 8^ followed by scRNA-seq of the freshly sorted live cells using the droplet-based 10X Genomics Chromium platform. Further details on the demographic and clinical characteristics of our donors, as well as details regarding our cell hashing strategy, may be found in **Table S1**.

After rigorous pre-processing, we retained 225,382 individual transcriptomes from 74 donors. As the samples in our dataset encompassed a broad set of disease conditions, brain regions, and chemistry versions for the 10X platform, we applied algorithms for batch correction and normalization with the goal of identifying microglial states that are conserved across conditions. To separate microglia from non-microglial populations, we used Seurat^23^ to perform Louvain clustering, choosing a resolution where distinct cell types can be identified by canonical markers. In this initial clustering (**Figure S1**), small numbers (<5%) of adaptive immune cells, monocytes, erythrocytes, and other non-immune populations segregated from our microglia and were not further analyzed.

### Microglial subpopulations and signature genes

We then subclustered myeloid/microglial cells to derive a shared reference model across all regions and diseases. After selecting a model where all pairs of clusters had less than 20% ambiguous assignment of cells using multilayer perceptron classification **(Methods)** and retaining clusters with >100 cells, we prioritized a population structure consisting of 12 distinct clusters (**Figure 1A**). The mean number of unique molecular identifiers (UMIs) and genes detected in microglia is similar across batches, technologies, and clusters (**Figure S2A-F**, **Table S1**), and *post-hoc* computational cluster validation supported the stability of this cluster structure (**Figure S2G)**. We first confirmed the microglial identity of our clusters by evaluating a set of core microglial genes^5, 24–26^ as well as monocyte (*S100A8*, *VCAN*) and macrophage (*SELL*, *EMILIN2*, *GDA*) genes (**Figure 1B**). Notably, all 12 clusters express *AIF1* and *C1QA*, well-validated markers of microglial identity in the brain, but some microglia-specific murine marker genes, such as *HEXB*^27^, are expressed at low levels or are inconsistent in our human data. We also examined proposed markers (*LYVE1*, *MS4A7*, and *CD163*) of the border-associated macrophage (BAM) subset recently reported in mice^28, 29^, and no cluster appears to be predominantly composed of BAM-like cells.

**Figure 1.**
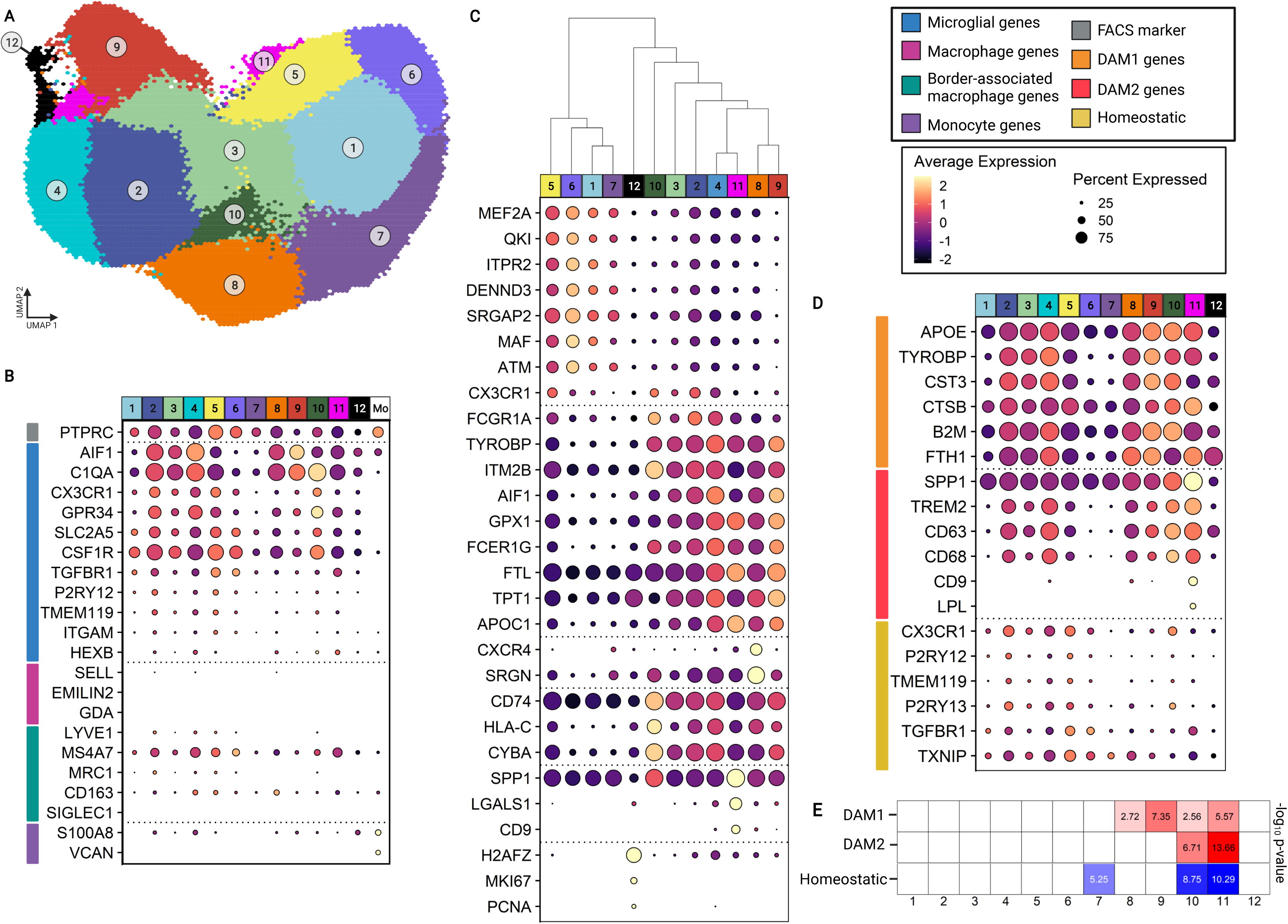
Microglial subtypes are defined by distinct marker genes and shared expression programs. **(A) Visual representation of the 12 microglial subtypes.** A hex-binned uniform manifold approximation and projection plot presents microglial subsets: other cells are shown in **Figure S1**. Each hexagon is colored by the majority cluster identity among all cells aggregated (mean of 50 cells per hexagon). **(B) Expression levels of genes delineating different myeloid identities.** The box over panel D summarizes the selected gene sets, which are color-coded on the left side of each panel. In panel B and each following panel, each column presents data from a cluster of cells (microglial subtypes colored as in **A** and monocytes, Mo), and each row represents the level of expression of a gene. The size of the circle represents the percentage of cells in each cluster that express the gene. The color of the circle represents z-scored gene expression. Genes were chosen for association with microglial, macrophage, border-associated macrophage, or monocytic identity. **(C) Subtype-enriched marker genes.** Marker genes, selected by pairwise differential expression testing with MAST, delineate broad microglial families with overlapping gene expression programs and small clusters with strongly distinguishing marker genes. Hierarchical clustering with complete linkage on the expression of genes shown in this figure is shown by the dendrogram at the superior aspect of the figure. (**D) Expression level of DAM gene sets and homeostatic genes across microglial subsets. (E) Heatmap of DAM gene set enrichment.** Enrichment of DAM subtype signature genes in upregulated (for DAM1/DAM2, in red) or downregulated (homeostatic, in blue) genes associated with each cluster is shown. Each column is one microglial subtype. Enrichment was tested by false discovery rate-corrected hypergeometric test. DAM disease-associated microglia. See also **Table S2**.

Next, we performed pairwise differential expression analysis to define the genes that best differentiated our microglial subtypes (**Methods, Table S2**). Representative distinguishing genes are shown in **Figure 1C**. The clusters are numbered in descending order based on their size, with 1 having the largest number of cells and 12 the least. Clusters 1, 2, and 3 are the most abundant clusters in most individuals (**Figure S3**), and cluster 3 shows marker expression suggesting that it is intermediate between clusters 1 and 2. Notably, clusters 2 and 3 are more enriched in genes associated with classical homeostatic-active states^5^ (*CX3CR1, FCGR1A, P2RY12*), and these two clusters have relatively few differentially upregulated genes compared to all other microglial clusters. On the other hand, genes upregulated in cluster 1 (the largest cluster) include disease genes such as *ITPR2* and *SORL1*, as well as transcription factors and RNA-binding proteins, such as *MEF2A*, *RUNX1*, and *CELF1*. This suggests that clusters 2 and 3 may be closest to the classic description of “homeostatic” microglia, while cluster 1 and its closely related family are a divergent branch of microglial differentiation. Cluster 6 is the closest transcriptionally to cluster 1, expressing high levels of *SRGAP2* and *QKI*, an RNA-binding protein that regulates microglial phagocytosis in the context of demyelination^30, 31^. Interestingly, cluster 5 appears to be an alternate intermediary state between clusters 1 and 2, as it expresses *CX3CR1* alongside *QKI* and *MEF2A*. In contrast, clusters 4 and 9, which are transcriptionally adjacent to cluster 2, have an overlapping set of enriched genes, including *C1QA*, *TYROBP*, *ITM2B*, *GPX1*, and *FCER1G*. Thus, the broadest division in microglial subtypes appears to be between clusters 2/4/9 on the left side of our cluster structure, and 1/5/6/7 on the right side of the cluster structure. Clusters 8 and 10 are located between these broad families but are more homologous to 2/4/9. Cluster 8 is enriched in *CXCR4* and *SRGN*, while cluster 10 is enriched in *HLA*-*C, CD74* (two proteins that play important roles in antigen presentation), and *CYBA*. Cluster 11 also shares some transcriptomic homology with 2/4/9 but is distinguished by enrichment in *SPP1* and *LGALS1*. Finally, cluster 12 expresses *MKI67* and *PCNA*, suggesting a proliferative phenotype.

The disease-associated microglial (DAM) state^6^ has been clearly defined in mouse models, but results in human studies have been mixed^4, 16, 32, 33^. The lack of clarity around the possible presence and/or role of DAM genes in humans may stem from technical differences between studies and the relatively small numbers of microglia profiled in studies to date. In addition, the proposed transition from homeostatic to DAM1, an initial TREM2-independent state, then to DAM2, a later, TREM2-dependent state, has been incompletely explored in humans. We reasoned that separately examining the enrichment of signatures associated with both DAM sub-states might allow us to delineate the distribution of human microglia along this proposed DAM trajectory. Markers for both DAM subsets, as well as homeostatic genes downregulated in the DAM progression, are shown in **Figure 1D**. We hypothesized that, if there was a DAM subset in our dataset, it would likely be a small, distinct subset primarily enriched in the DAM2 signature due to the predominance of autopsy tissue from late-stage neurodegenerative disease among our data. Indeed, we identified a small subpopulation, cluster 11, representing 1% of our microglia, that shows strong enrichment for the DAM2 signature. However, as seen in **Figure 1E**, the situation is complex: The DAM signature genes are expressed in four different microglial clusters showing different combinations of DAM1 and DAM2 enrichment, with the DAM1 signature being most enriched in cluster 9. We note that cluster 10, the *CD74*^high^ cluster that we had highlighted in our prior report as DAM-enriched^4^, also shows significant enrichment in the DAM2 signature, albeit at lower level than cluster 11. Notably, clusters 10 and 11 both show substantial downregulation of the homeostatic microglial signature identified in the original DAM publication^6^, while cluster 9 does not demonstrate significant downregulation of the homeostatic gene set, suggesting that clusters 10 and 11 are further along the trajectory of divergence from the homeostatic microglial phenotype. Our data suggests that, in humans, there may be distinct microglial subtypes with different combinations of DAM-related transcriptional programs that fulfill different functions, rather than a single linear DAM trajectory.

### Axes of metabolic and functional variation across microglial subtypes

We then evaluated inter-cluster relatedness by using a post-hoc machine-learning approach leveraging a multilayer perceptron classifier to examine the homology of gene expression programs between cells assigned to different clusters^34^. We visualize these results in a constellation diagram (middle of **Figure 2A**), where the thickness of the line between clusters is proportional to the number of cells between the clusters that are misclassified, and the size of the nodes corresponds to the number of cells in that given cluster. A central question in microglial biology is how different subtypes may branch off from core homeostatic phenotypes, and whether trajectories of microglial state transition are linear or characterized by critical bifurcation points. Based on our analysis, cluster 3 exhibited the most “central” gene expression profile. As expected, clusters 2 and 4 show substantial homology to one another, as do clusters 1 and 6. Notably, cluster 5, another prospective intermediate state, shows overlap with both clusters 1 and 2, although it is more homologous to cluster 1. This highlights a continuous transition between extremes of gene expression in either direction along this central division. To explore the functional relevance of this division, we used topGO to conduct Gene Ontology (GO)^35–, 37^ analysis on top differentially expressed genes in each cluster and summarized results with rrvgo^38^. As shown on the left and right sides of **Figure 2A**, the most heavily enriched terms along the left (clusters 4,9) side of our population structure are related to metabolism, particularly oxidative phosphorylation, catabolism, and peptide metabolism, and immune response and localization. In contrast, the right side (clusters 1,6) of our population structure shows enrichment of alternative metabolic pathways, including heterocyclic metabolism and nitrogen-containing compound metabolism as well as transcriptional regulation. Intriguingly, the intermediate cluster 5 shows strong association with motility (**Figure S4A).** This highlights a central divide in metabolism and function, suggesting a homeostatic-active phenotype on the left side of our model that transitions to different metabolic and functional phenotypes on the right side of our structure, with intermediate states that may play different functional roles. Cluster 9, which has a partially overlapping transcriptional signature with cluster 4, is the most closely related to cluster 12, which shows enrichment for proliferation and oxidative phosphorylation (**Figure S4B)**. Cluster 9 also shows substantial transcriptomic similarity to cluster 11, which is enriched for lipid processing and beta-amyloid clearance (**Figure S4C**), consistent with the proposed role of TREM2 in this signature^33^, confirming the continuum of DAM transitional states identified in our earlier analysis (**Figure 1E**).

**Figure 2.**
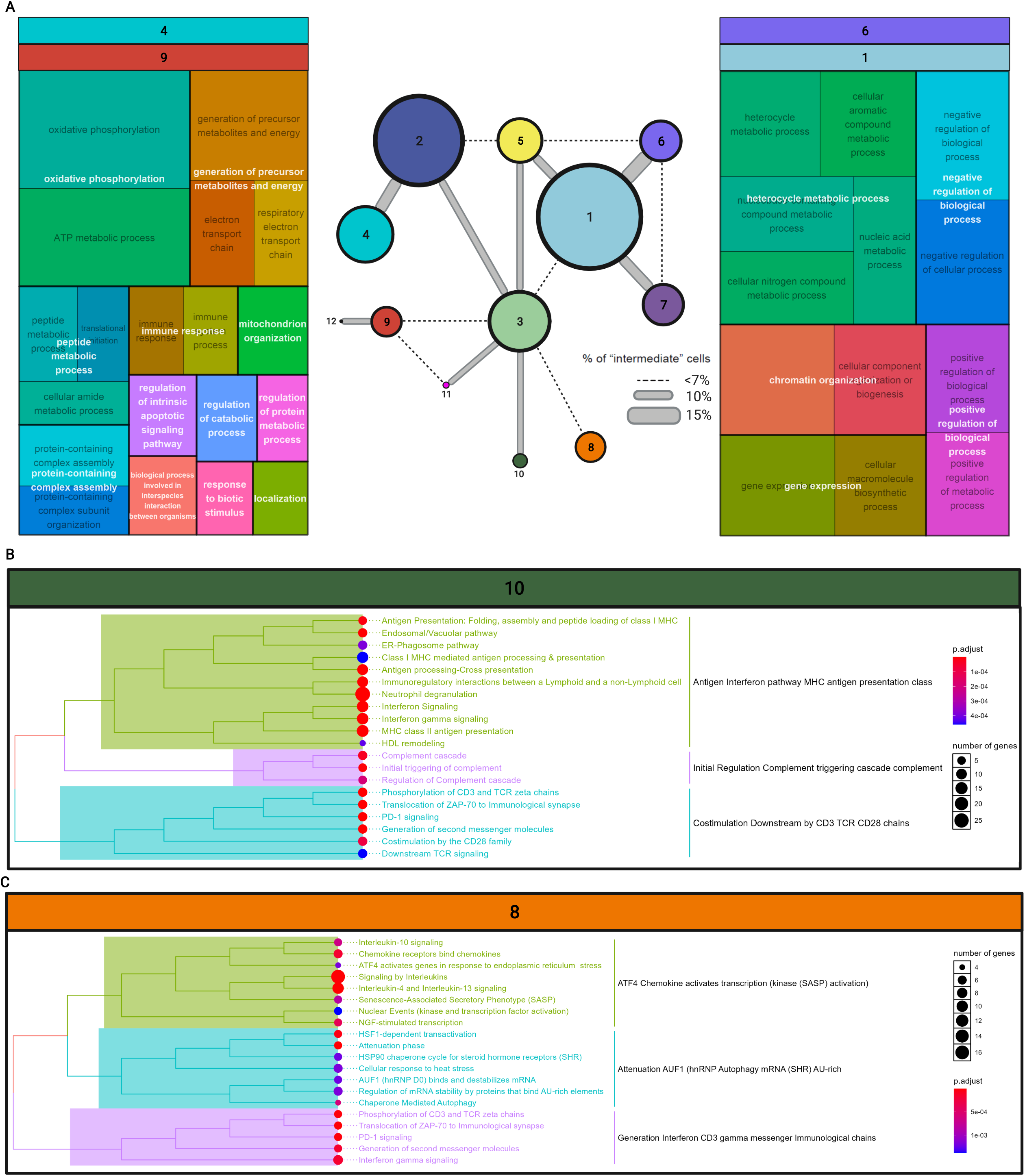
Microglia display a complex trajectory of state transition with several primary axes. **(A) A central metabolic divide separates divergent subtype families.** Constellation diagram demonstrates relationships between clusters by way of post-hoc classification. Each pair of distinct clusters was used to train a multilayer perceptron 50 times using 5-fold cross validation to obtain a classification for every cell. Cells that were classified to the same cluster less than 40 times were considered ambiguous. The fraction of ambiguous cells determines the width of the connecting lines in the diagram. Each node is a single cluster, with size scaled in proportion to the number of cells contained therein. Notably, even closely related clusters can be reliably distinguished over 85% of the time. Cluster 3, which has few distinct marker genes, has the most “central” expression profile, with close relationships to the 2/4 family and the 1/6/7 family. Cluster 5 represents another intermediate step between the 2/4 and 1/6/7 families. Gene ontology (GO) annotation was performed with topGO and summarized with rrvgo. Parent terms are shown in white, overlaid over child terms. GO annotation for clusters 1/6 and clusters 4/9 reveals a metabolic shift between the two groups: clusters 4/9 show enrichment of oxidative phosphorylation, catabolism, and protein metabolism, as well as general immune response, while clusters 1/6 demonstrate upregulation of heterocyclic and nitrogen-containing compound metabolism alongside transcriptional regulation. **(B) Clusters 8 and 10 share a signature of interferon-gamma signaling and antigen presentation but differ in other pathways.** Reactome annotation of clusters 8 and 10 aggregated by group highlights shared enrichment for T-cell interaction and interferon gamma signaling (purple in cluster 8 and blue in cluster 10). Cluster 10 shows upregulation of complement signaling (purple) and MHCI/II antigen presentation (green), while cluster 8 shows upregulation of chaperone and steroid signaling (blue) and interleukin signaling (green). See also **Figure S4**.

Clusters 8 and 10, whose signatures suggest substantial microglial activation, are clearly distinguishable from other clusters, although they maintain a relationship to the central cluster 3. To explore this axis, we annotated clusters using ClusterProfiler to perform Reactome^39–41^ pathway analysis (**Figure 2B/C)**. Some overlap was present between these two clusters, as both clusters contain genes associated with antigen presentation, interferon signaling, and T-cell interaction. However, cluster 10 exhibits stronger association with both class I and class II MHC signaling and complement signaling. In contrast, cluster 8, which expresses significant levels of early response genes, shows upregulation of pathways associated with chaperone signaling, steroid response, interleukin signaling (particularly *IL4*/*IL10*/*IL13*), and the senescence-associated secretory phenotype. These phenotypic differences can also be recapitulated with GO annotation (**Figure S4D**).

Thus, this analysis highlights the divergent nature of the microglial differentiation program, suggesting that there are at least 3 distinct tracks of microglial subtype specification that emerge from the most basal microglial state, including a central metabolic divide, an axis of immunological activation, and a trajectory that contains elements of the DAM signature identified in murine model systems. These tracks appear to be nonlinear and different paths of transition may exist between terminal states, a result that is consistent with an ancillary pseudotime analysis leveraging the Monocle3^42–44^ algorithm that defines a complex trajectory (**Figure S4E**). In addition, consistent representation of clusters across donors, regions, and diseases (**Figure S3A/B**) supports the conceptual framework of our trajectory analyses, as this shared representation supports the idea that we are examining an actual biological continuity rather than identifying state shifts that result from differences in disease or region. Notably, the results of our analyses further underscore the robust nature of our population structure, as even clusters with substantial overlap of gene expression signatures are still robustly differentiable more than 85% of the time. Further work *in vitro* and *in situ* will be required to confirm our observations and to understand the importance of functional and metabolic shifts in health and disease.

### Microglial subset proportions across regions and diseases

To illustrate the nature of our data, we report the proportion of different microglial subtypes in each disease, region, and individual. (**Figure 3**, **Figure S3**). First, we note that most clusters are present in each individual, albeit at different frequencies (**Figure S3A**). The most common microglial subtypes in most individuals are clusters 1 to 6, suggesting that these subtypes capture a homeostatic spectrum and that neurodegenerative diseases involve small shifts towards distinct microglial states. Notably, grouping samples by region (**Figure 3A/B**) or diagnosis (**Figure 3C/D**), demonstrates the fact that even with different numbers of cells from different regions and diagnoses, we see a similar distribution of populations across diseases and regions. Much larger datasets will be needed to directly identify disease associations.

**Figure 3.**
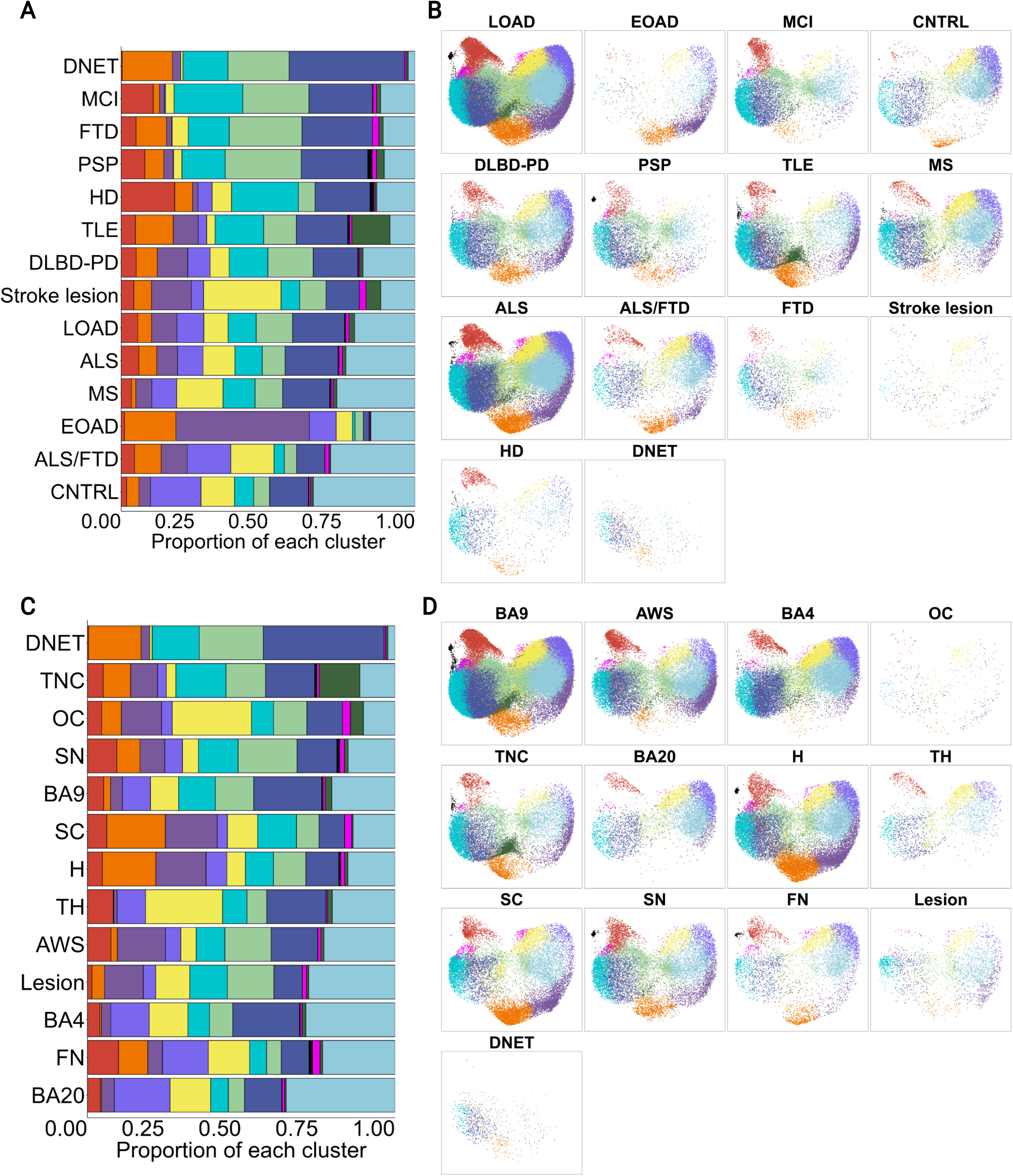
Human microglial subsets are found across diseases and regions. **(A)-(D) Microglial subsets are broadly represented across diseases and regions.** On the left (A, C), each bar shows the proportion of each cluster among all microglia from a given disease. On the right (B, D), UMAP plots are split by disease. Plots are color-coded in accordance with Figure 1A. Most subsets are represented across all diseases and all regions, albeit in different numbers. LOAD late-onset Alzheimer’s disease, EOAD early onset Alzheimer’s disease, MCI mild cognitive impairment, CNTRL control, DLBD-PD diffuse Lewy body disease-Parkinson’s disease, PSP progressive supranuclear palsy, TLE temporal lobe epilepsy, MS multiple sclerosis, ALS amyotrophic lateral sclerosis, FTD frontotemporal dementia, HD Huntington’s disease, DNET dysembryoplastic neuroepithelial tumor, BA Brodmann area, AWS anterior watershed, OC occipital cortex, TNC temporal neocortex, H hippocampus, TH thalamus, SC spinal cord, SN substantia nigra, FN facial nucleus. See also **Figure S3**.

### Annotating disease and trait associations of distinct microglial subsets

Although direct testing of disease associations in our dataset is underpowered given the design of our study, the depth and quality of our sequencing data enabled us to pursue gene enrichment analyses to implicate certain subsets in different diseases. For MS, we utilized a recent publication from the International Multiple Sclerosis Genetics Consortium that identified a comprehensive set of 551 MS susceptibility genes^45^. We found that clusters on the right side of our microglial cloud, specifically clusters 5 and 6, were significantly enriched in MS susceptibility genes, highlighting a possible role of one arm of our microglial differentiation tree in MS susceptibility (**Figure 4A**). Next, we explored the genome-wide association study (GWAS) catalog^46^, a curated database containing SNP-trait associations from GWAS studies. As seen in **Figure 4B**, we recapitulated the enrichment of MS in cluster 6 in this database, although cluster 5 did not pass the threshold for significance in this analysis. Clusters 1 and 6 also show enrichment for other neurodegenerative and neuropsychiatric disease genes, including AD, PD, and depression. Similarly, cluster 10 shows enrichment for MS and schizophrenia. The strong complement expression found in cluster 10 aligns with previous reports of the role of complement-related genes in schizophrenia^47^. Finally, cluster 8, the *CXCR4*-enriched cluster, has a set of disease associations that suggest a role in conditions characterized by neuroinflammatory signaling but not neurodegenerative diseases. Interestingly, we found no substantial enrichment of disease genes associated with stroke, seizure, ALS/FTD, or glioma. This may speak to either a less important role of microglia in the primary pathogenesis of these diseases or less extensive GWAS annotation of these diseases.

**Figure 4.**
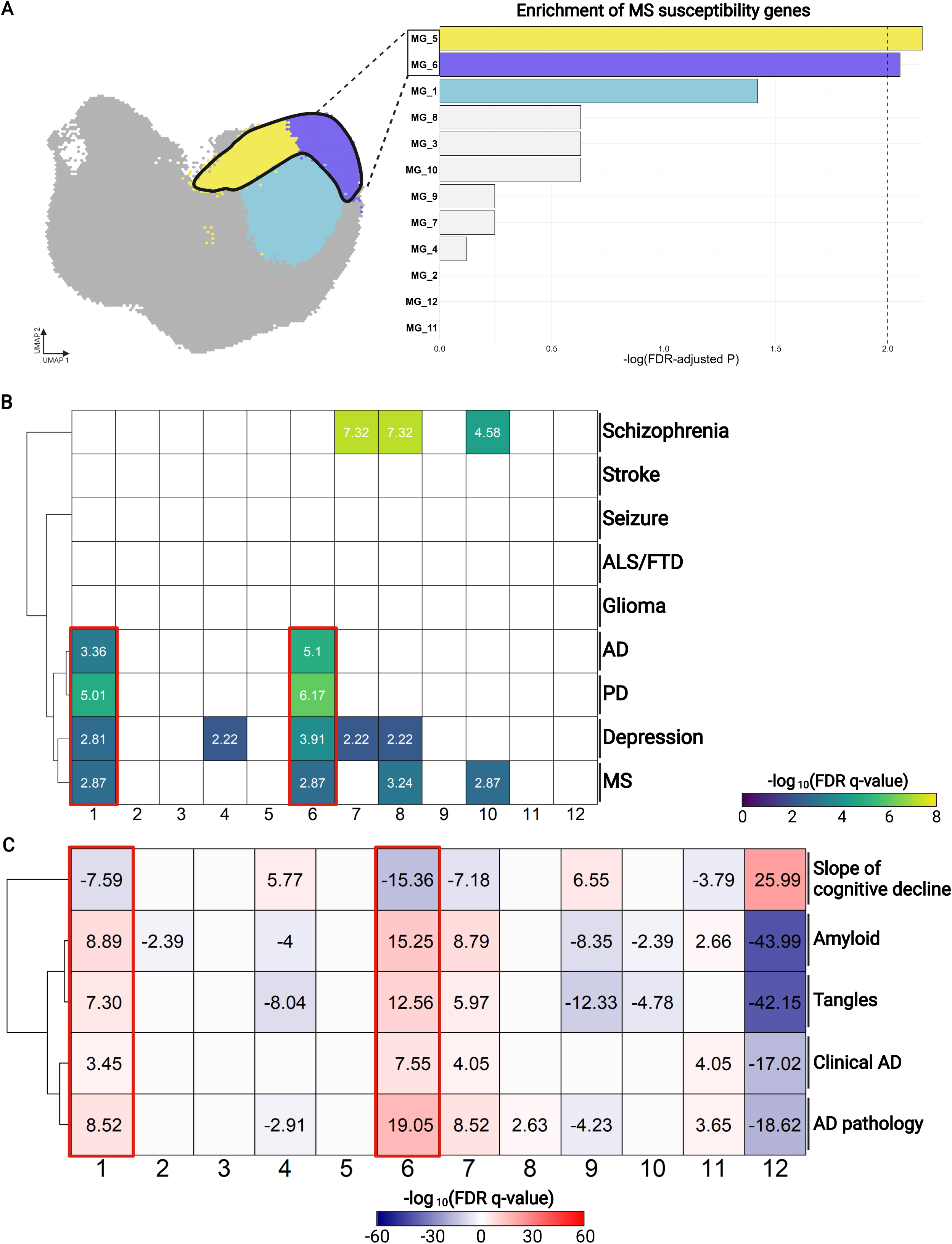
Disease annotation implicates specific microglial families in disease. **(A) Clusters 5 and 6 are enriched in GWAS-derived MS susceptibility genes.** The y-axis of the barplot shows the different clusters, ranked in descending order of the negative log-transformed p-values on the x axis. Enrichment of MS susceptibility genes in upregulated gene lists associated with each cluster was tested with the hypergeometric test with Benjamini-Hochberg correction. Bars are colored if they have an FDR < 0.01. **(B) Clusters 1 and 6 are enriched in genes associated with neurodegenerative diseases.** Enrichment analysis of genes associated with each disease in the GWAS catalog was performed with same parameters. Diseases are listed on the y-axis, and negative log-transformed p-values are shown for combinations of clusters and traits where they have an FDR < 0.01. Coloration of squares corresponds to p-value magnitude: larger p-values correspond to darker blue squares, whereas smaller p-values correspond to yellow coloration. **(C) Clusters 1 and 6 correlate with clinical and pathological traits in AD.** In this case, enrichment was performed separately for both the genes positively and negatively correlated with each trait in upregulated genes for each cluster. Coloration of each box relates to the strength and directionality of each association. Red (positive numbers) corresponds to genes upregulated with the trait, while blue corresponds to genes downregulated in relation to the trait. AD Alzheimer’s disease, PD Parkinson’s disease, MS multiple sclerosis, ALS amyotrophic lateral sclerosis, FTD frontotemporal dementia, GWAS genome-wide association study. See also **Table S3**.

To complement these analyses, we repeated our earlier analysis^4^ evaluating AD-related traits more thoroughly. We leveraged associations with traits from a large analysis of bulk cortical (BA9) RNA sequencing data from 1092 participants in the ROSMAP^48, 49^ cohorts (**Table S3**); these individuals do not overlap with the ROSMAP participants included in our microglial dataset. These bulk RNA sequencing data contain transcripts from all parenchymal cells, including microglia. To evaluate the enrichment of microglial clusters for genes associated with each trait, we evaluated the overlap of cluster-specific signature genes with gene sets that were significantly positively or negatively associated with each of these traits (**Figure 4C**). Intriguingly, clusters 1 and 6, the right side of the microglial cloud, are enriched for genes which are positively correlated with amyloid-beta pathology, tau tangle pathology, and both a clinical and pathological diagnosis of Alzheimer’s disease. Consistent with these results, they are also enriched for genes negatively correlated with the slope of cognitive decline where a larger negative number indicates worsening cognitive dysfunction. In contrast, clusters 2, 4, 9, and 10 are enriched in genes negatively correlated with tau and amyloid pathology, and clusters 4/9 are enriched in genes positively correlated with the slope of cognitive decline. Further, our DAM2^high^ cluster 11 is enriched for genes positively associated with AD and amyloid pathology, but not tau pathology. This cluster also shows enrichment for genes negatively correlated with cognitive decline. Our results strongly suggest that the AD cortex is enriched for genes defining clusters 1 and 6; this could occur either from an increase in the proportion of these subtypes in AD cortical tissue or from the enhanced expression of these signature genes in the microglia of the AD cortex. It also implicates cluster 11 more modestly, suggesting a narrower contribution to amyloid rather than tau proteinopathy. Notably, several molecular pathologies that we evaluated - including cerebral amyloid angiopathy, arteriolar sclerosis, cerebrovascular disease, and TDP-43 pathology - showed no enrichment with any of our clusters. Interestingly, cluster 12 showed enrichment trends akin to clusters 2, 4, 9, and 10, perhaps because of the heavy overrepresentation of genes associated with oxidative phosphorylation. Overall, by leveraging indirect disease annotation, we identify a family of microglial subtypes that are strongly enriched in genes associated with neurodegenerative disease and associated with AD traits in an independent ROSMAP cohort.

### A robust co-detection workflow leveraging immunofluorescence and RNAscope recapitulates microglial subset abundances *in situ*

Having identified microglial subtypes, we sought to validate their existence *in situ*. We had previously done this with immunofluorescence^4, 10^, but the range of potential antibody markers is limited and does not necessarily reflect RNA expression. Thus, we developed a co-detection workflow that merged *IBA1* staining (an ubiquitous marker of myeloid cells in the brain at the protein level), with Advanced Cell Diagnostic’s RNAscope protocol^50^ for fluorescent *in situ* hybridization to allow for single-molecule RNA detection. We coupled this experimental pipeline to CellProfiler v4.2.1^51–54^ for automated segmentation of image data (**Methods**). This workflow enables the capture of microglia-specific gene expression, localization of transcripts within microglia, and structured assessment of cellular morphology.

To illustrate our approach, we chose two panels of genes to discriminate different microglial subsets *in situ*. In panel 1 (**Figure 5A-D, Figure S5A**), we used probes against transcripts of *CD74*, a gene that we found to be relatively enriched in immunologically active subtypes (clusters 2, 3, 4, 8, 9, and 10) and downregulated in clusters 1,5, and 6. We have previously reported the existence and relevance of a *CD74^high^*microglial subset^4^, and the equivalent subset in our current model (cluster 10) has a 1.5 fold greater expression of CD74 relative to all other clusters (**Figure 5A**). We also included probes against *CXCR4*, a gene predominantly expressed in cluster 8. This panel separates the 1/5/6 family from other clusters, exploring our primary axis of variation, and allows for the discrimination of the two subsets (8 and 10) in a second axis of variation associated with antigen presentation and immune cell interaction. In panel 2 **(Figure 5E-H, Figure S5B)**, we included probes against *GPX1*, a gene predominantly expressed in clusters 4, 9, and 11, as well as *SPP1*, a previously proposed DAM marker that is enriched in cluster 11. This panel enables a more detailed examination of clusters along the DAM-like trajectory that were not captured in the first panel. We stained tissue sections from individuals with pathological diagnoses of AD, PD, PSP, or DLBD and individuals with age-related tauopathy (**Table S4**).

**Figure 5.**
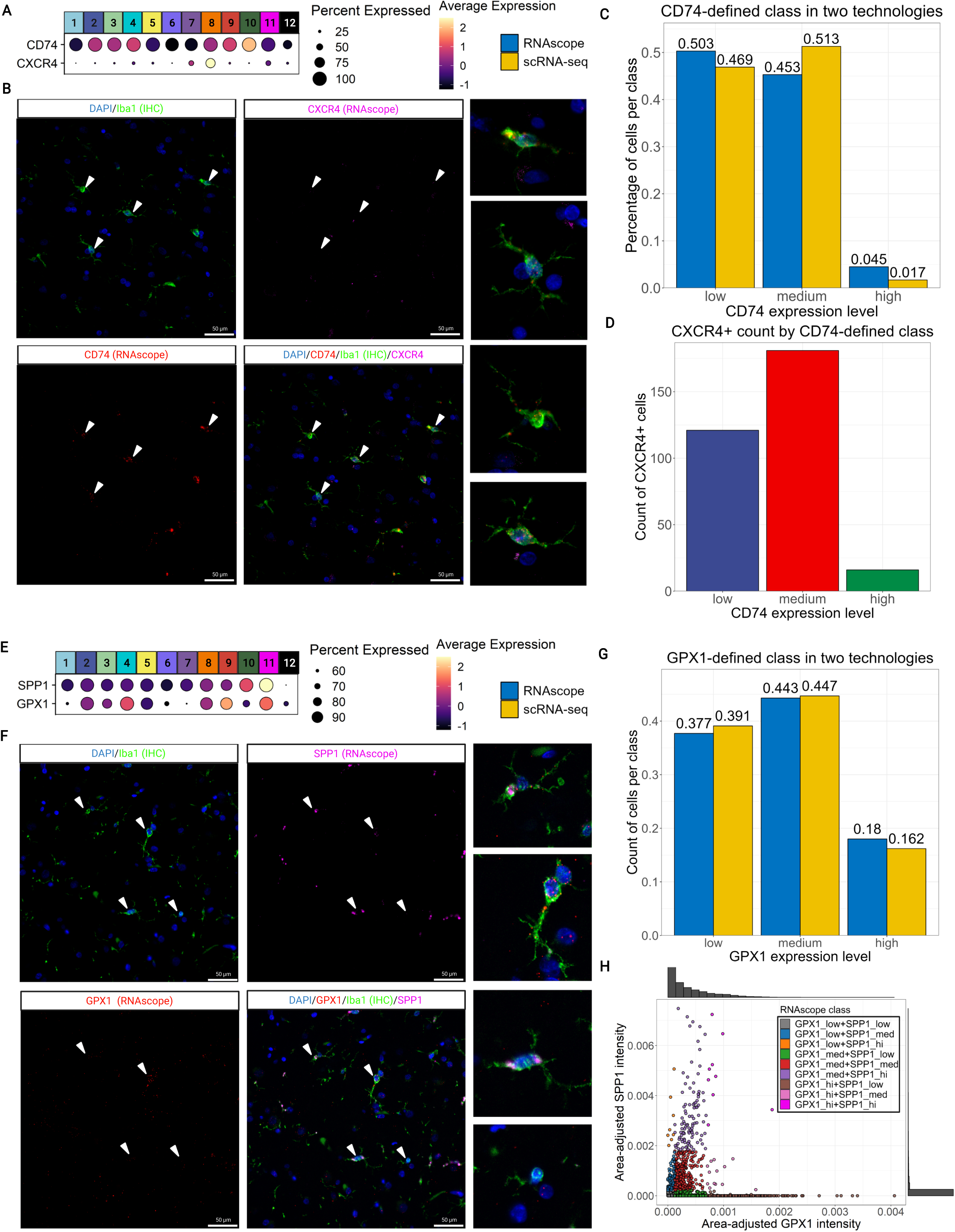
*In situ* confirmation of microglial population structure with joint immunofluorescence-RNAscope with automated segmentation. **(A) *CD74* demarcates a small, immunologically active subset, while *CXCR4* delineates a distinct immunologically active subset.** The size of the circle represents the percentage of cells in each cluster that express the gene, and the color of the circle represents z-scored gene expression. *CD74* is overexpressed in cluster 10, while *CXCR4* is primarily expressed in cluster 8. **(B) Representative images showing *CD74* and *CXCR4* in IBA1^+^ microglia.** RNAscope staining for *CD74* (red) and *CXCR4* (pink) in IBA1^+^ microglial cells (green) in human cortical brain slices, with nuclear DAPI staining (blue). In the same field of view, microglia with different levels of *CD74* and with or without expression of *CXCR4* can be observed (arrowheads point to representative microglia). **(C) Separating single *in situ* cells using *CD74* expression thresholds adapted from scRNA-seq identifies similar proportions across technologies.** The proportion of cells, along the y-axis, that express low, medium, or high levels of *CD74*, along the x-axis, in scRNA-seq is shown in yellow, while *in situ* results (area-adjusted *CD74* expression binned on thresholds from scRNA-seq) are shown in blue. **(D) *CXCR4*^+^ cells match expected distribution within *CD74* expression classes.** *CD74* expression class, as described in (**C**), is shown on the x-axis, and count of CXCR4^+^ cells is shown on the y-axis. CXCR4^+^ microglial cells are identified *in situ* and most fall into the CD74^med^ class, confirming our scRNA-seq findings. **(E) *GPX1* and *SPP1* delineate the DAM axis and extremes in homeostatic-active families. (F) Representative images from our joint staining protocol for *GPX1* and *SPP1*.** Staining as in (B), except that RNAScope *SPP1* is pink and *GPX1* is red. **(G) Separating single *in situ* cells on the basis of *GPX1* expression thresholds borrowed from scRNA-seq also identifies similar proportions across technologies.** Analysis performed as in (C) but using *GPX1* expression data. **(H) Gradated expression of both *SPP1* and *GPX1*.** Individual cells are plotted as single dots, where the axes represent area-adjusted expression of *GPX1* (x) or *SPP1* (y). DAM disease-associated microglia. See also **Figure S5** and **Tables S4/5**.

Representative images for panel 1 are shown in **Figure 5B**, demonstrating the capture of microglia-specific transcripts, including those in the distal processes of single microglia. By taking area-normalized *CD74* expression per cell and binning into low expression, medium expression and high expression based on fold change thresholds derived from our single-cell data (**Methods**), we found that the proportions of cells in each bin are similar to the proportions of cells in our single-cell dataset (**Figure 5C**). The *CD74*-low grouping includes clusters 1, 5, 6, 7, and 12, *CD74*-medium represents clusters 2, 3, 4, 8, 9, and 11, and *CD74*-high identifies cluster 10. Moreover, concurrently examining expression of *CXCR4* across all 3 bins demonstrates peak *CXCR4* expression in the *CD74*-medium populations, in agreement with our scRNA-seq data (**Figure 5D**). Thus, panel 1 validates clusters 8 and 10 *in situ* and demonstrates our ability to discriminate multiple distinct microglial subsets in each field of view. As shown **in Figure 5E/F**, panel 2 was designed to discriminate the homeostatic-active family and concurrently identify clusters on the DAM axis, particularly cluster 11, that our first panel cannot capture. We first evaluated *GPX1*, and, as with panel 1 markers, we observe similar proportions of cells in the different *GPX1* categories between the *in situ* and scRNAseq approaches, confirming the translatability of our scRNAseq-derived cluster definitions (**Figure 5G**). Notably, we identify cells co-expressing different levels of *GPX1* and *SPP1* and subpopulations that appear similar to clusters 9 and 11, making them good markers for future investigations targeting the DAM-like axis (**Figure 5H**). This second panel thus offers an independent validation of a different aspect of our population structure, providing markers for future study design.

To begin to explore cell shape across subtypes, we evaluated morphological measures captured from each microglia by CellProfiler. First, *CD74*, *SPP1*, and *GPX1*-medium classes have the highest level of compactness, a measure where higher values indicate more ramification, suggesting maximal ramification in these partially overlapping subtypes (**Figure S5C**). Similarly, eccentricity - a feature where 0 denotes a perfect circle and 1 denotes a line - is lowest in both the CD74^high^ and GPX1^high^ classes and highest in the CD74^low^ and GPX1^low^ classes (**Figure S5D**), suggesting that the latter subtypes have a more elongated morphology. Microglial activation induces process retraction and an amoeboid morphology^55^, and in our scRNA-seq data, cells expressing high levels of CD74 and GPX1 express markers of activation. Interestingly, CXCR4^+^ cells exhibit higher CD74 radial distance on average (**Figure S5E)** and higher compactness scores (**Figure S5F**), suggesting that, unlike other activated classes, CXCR4^+^ cells are likely to be particularly ramified.

We therefore validate our cross-disease scRNA-seq resource by identifying the derived clusters *in situ* and exploring the morphology of different subtypes. We also highlight the existence of cluster 8 cells in the tissue, supporting our hypothesis that the stress response observed in certain *ex vivo* preparations of microglia is a natural transcriptional program, albeit one that may be induced by certain microglial manipulations^56, 57^.

### Extending the use of our dataset as a reference: *in vitro* model systems, single-nucleus data, and other diseases

This data resource, which was designed to identify a shared, stable microglial population structure across diseases and regions, can be used to annotate other microglial datasets and to evaluate how well model systems recapitulate the microglial diversity observed in human tissue. We chose two datasets: a single-nucleus RNAseq dataset (see **Methods**) from the ROSMAP^48, 49^ cohorts of older individuals with and without AD and a single-cell myeloid dataset^58^ from surgical resections of tissue from glioblastoma, a disease where microglia are thought to play a central role^59^. In parallel, we chose to explore two human iPSC-derived microglial datasets: a murine xenograft system^60, 61^ and an *in vitro* system used for CRISPR screening^62^. Label transfer was then performed for these four datasets. To simplify the classification task, transcriptionally similar groups (i.e., 2 and 4 or 1, 6, and 7) in the reference data were grouped. We then applied a pipeline leveraging consensus voting of pairwise support vector machine (SVM) classifiers to classify the smaller, more transcriptionally unique subtypes and a flat XGBoost^63^ (XGB) classifier retaining only classifications with confidence of 50% or higher for the larger clusters (**Figure 6A**). Quality metrics for our label transfer pipeline are shown in **Figure S6** and **Table S6**. In short, our approach is sensitive and specific, with joint accuracy averaging 85% across all models, and the lowest confidence assignments coming from clusters with inherently fluid boundaries, validating the efficacy of this pipeline.

**Figure 6.**
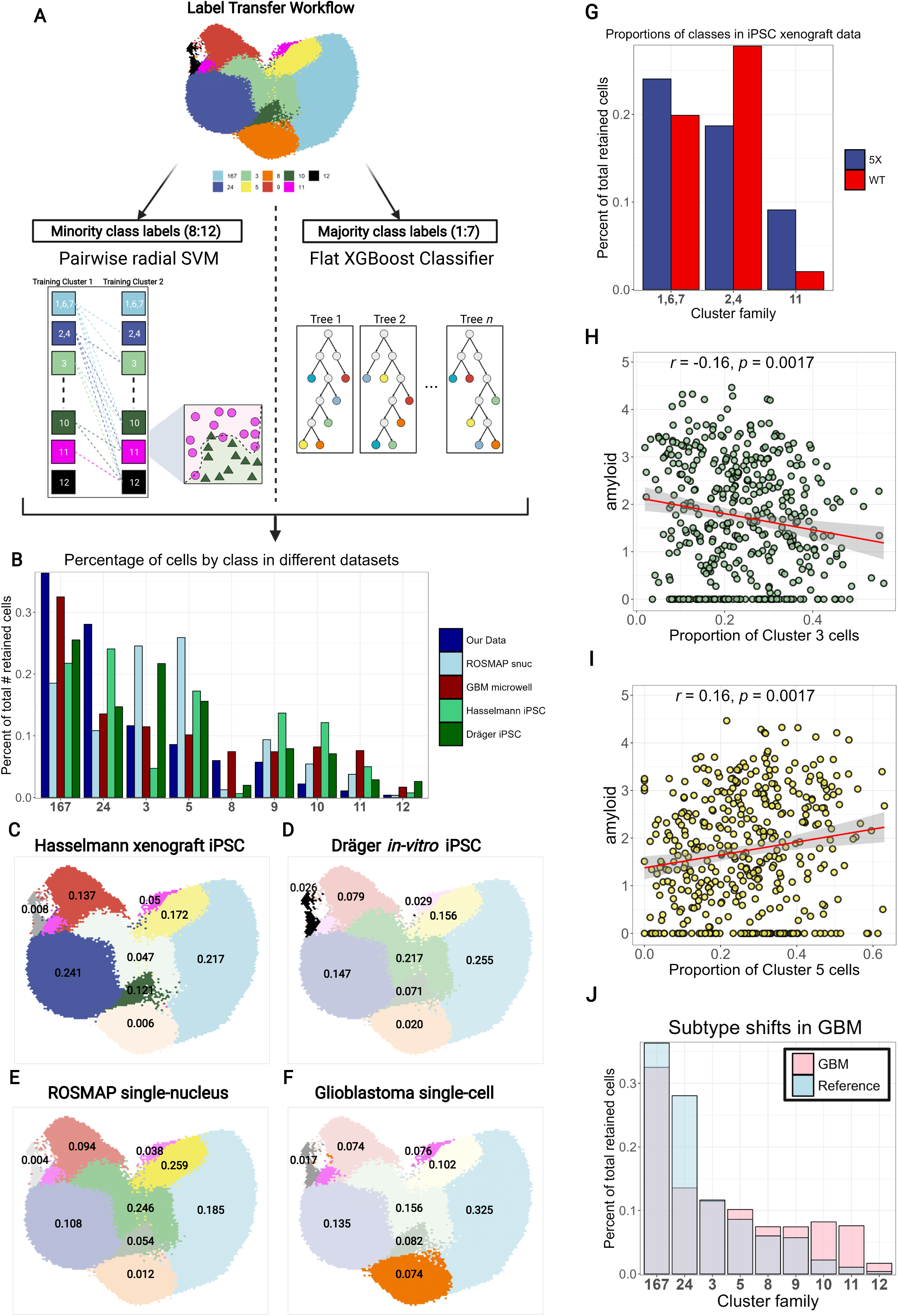
Live microglial population structure enables annotation of datasets from model systems and data produced with different technologies. **(A) Overview of our label transfer workflow.** Similar classes were aggregated (2 and 4 or 1, 6, and 7) to simplify the classification problem, and classifications from two types of models were merged to assign final class labels for all cells in query data. **(B) Distribution of subset proportions across different datasets in comparison to our reference. (C)-(F) Mapping of query datasets onto our reference model.** UMAP colors for each cluster family were shaded by the proportion of cells assigned to each family in each dataset. Numbers are the proportion of cells in each query dataset that were assigned to each cluster. **(G) Xenografted human iPSC microglia shift away from homeostatic-active phenotypes and towards disease-associated phenotypes in 5XFAD mice.** Barplot showing the proportion of iPSC microglia (y-axis) in each of 3 cluster families (x-axis) from either 5X (blue) or WT (red) mice. N=2 per condition. **(H) GBM induces depletion of homeostatic myeloid cells and shifts towards more inflammatory subtypes.** Barplot showing proportions of cells per group from the reference (blue), or the classified glioblastoma (GBM) data (red). Between the two datasets, the higher proportion is shown in its corresponding color, and the lower proportion is delineated in grey. **(H) Cluster 3 abundance correlates negatively with amyloid pathology in ROSMAP single-nucleus data.** Dotplot where each dot is a single donor. Axes are amyloid burden (y) and proportion of cells classified as cluster 5 (x). **(H) Conversely, cluster 5 abundance correlates positively with amyloid pathology.** See also **Figure S6** and **Table S6**.

Examining the percentages of cells classified into each cluster grouping across the different datasets (**Figure 6B-F**) demonstrates interesting differences. First, in both iPSC datasets (**Figure 6C/D)**, cells map to most of the states that we identified in our purified microglial dataset, suggesting that these model systems recapitulate a substantial amount of human *in vivo* heterogeneity. However, there are differences between the two contexts. The tissue-derived data (**Figure 6C**) shows higher fractions of microglia at terminal points in our axes of differentiation, such as the DAM-like axis of clusters 9, 10 and 11. Notably, we see a trend towards increased levels of cluster 11 in the 5X FAD mice, as well as a trend suggesting a shift from the 2/4 family to the 1/6/7 family in 5XFAD (**Figure 6G**), consistent with our AD enrichment analyses (**Figure 4B/C**). In contrast, the *in vitro* iPSC dataset contains higher numbers of proliferating cells (**Figure 6D**), “intermediate” cluster 3 cells, and inflammatory cluster 8 cells. These results highlight the utility of both model systems for modeling microglial diversity.

In turn, annotation of ROSMAP single-nucleus RNA-seq (snRNA-seq) data (see **Methods**) shows prominent representation of cells from the intermediate clusters 3 and 5 (**Figure 6E)**. Although snRNA-seq can be applied to frozen human brain tissue, it does not capture the cytoplasmic compartment^19, 20^. Thus, as the microglial subtype families 2/4 and 1/6/7 represent more polarized and differentiated branches of the homeostatic trajectory, albeit in distinct directions, technical differences may impair our ability to resolve some of the more distal phenotypes on our differentiation trajectory in nuclear data. Nonetheless, cluster 5, part of the same family that exhibits association with AD pathology (**Figure 4C**), shows a positive association with amyloid burden (**Figure 6H**), while cluster 3 shows a negative association with the same trait (**Figure 6I**) in the ROSMAP single-nucleus RNAseq-data. Finally, annotation of the glioblastoma dataset reveals high numbers of cells that map to the proliferative cluster 12, DAM2^high^ cluster 11, and cluster 8 (**Figure 6F**). Upregulation of SPP1, a gene defining cluster 11, has previously been reported in GBM-associated myeloid cells and has been shown to correlate with worse survival in human GBM patients^64^. Comparison of subtype proportions with our reference dataset from neurodegenerative diseases reinforces the presence of a dramatic shift away from the core homeostatic gradient and towards more inflammatory myeloid subtypes in GBM (**Figure 6J)**. This is consistent with prior observations reporting shifts away from homeostatic phenotypes among myeloid cells found in the glioma microenvironment^59^.

We thus demonstrate the utility of our dataset for annotating datasets from a wide variety of sources, such as diseases not captured in our dataset, single-nucleus data, and human iPSC-derived microglial model systems. We identify shifts in phenotype that accord with those previously reported in GBM and demonstrate that iPSC datasets, both *in vitro* and *in vivo,* capture an impressive amount of the microglial heterogeneity that we have identified in isolated, living human microglia.

### Prediction and validation of compounds driving cluster-specific transcriptional signatures and subtype recapitulation *in vitro*

Next, we sought to leverage our data to understand how to use chemical perturbation to direct state transitions towards specific subtypes. This would enhance (1) *in vitro* modeling using iMG or monocyte-derived microglia-like cells^65^ and (2) drug discovery as such tool compounds could provide leads for therapeutic development of *in vivo* modulation. Here, we used the V1 database of the Connectivity Map^21, 22^ (CMAP), a dataset that contains transcriptomic data associated with thousands of chemical perturbations across a wide array of cell lines. To increase the power of our initial analysis, we grouped related microglial subtypes together, querying CMAP using RNA signatures for three groups of microglial subtypes: Clusters 1/6, Clusters 2/4/9, and Clusters 8/10, chosen because they capture two of the primary axes of variation among our microglial subtypes. An overview of our workflow is shown in **Figure S7A**. From our *in silico* CMAP analysis (representative example: **Figure 7A, Table S7**), we prioritized 14 compounds for validation. For our initial screen, we exposed the Human Microglial Cell 3 (HMC3) line^66^ to each of the 14 compounds guided by the dosage used in CMAP. Since HMC3 cells were not used in CMAP, we optimized concentrations of each compound to minimize effects on survival and morphology. Then, we tested the effects of each compound after 6 and 24hr treatment by assessing the expression levels of two selected marker genes for each of the three groups of microglial subtypes using RT-qPCR, repeating this experiment at least three times with different batches of HMC3 cells (**Figure 7B-D** and **Figure S8B-D**). Four compounds met our pre-determined criteria for the screen: Torin-2 and Narciclasine both drove upregulation of marker genes associated with clusters 1/6, while Camptothecin and PMA drove 8/10 markers. Our results for compounds associated with clusters 2, 4, and 9 were inconclusive, as the marker genes that we chose did not show significant upregulation with these compounds.

**Figure 7.**
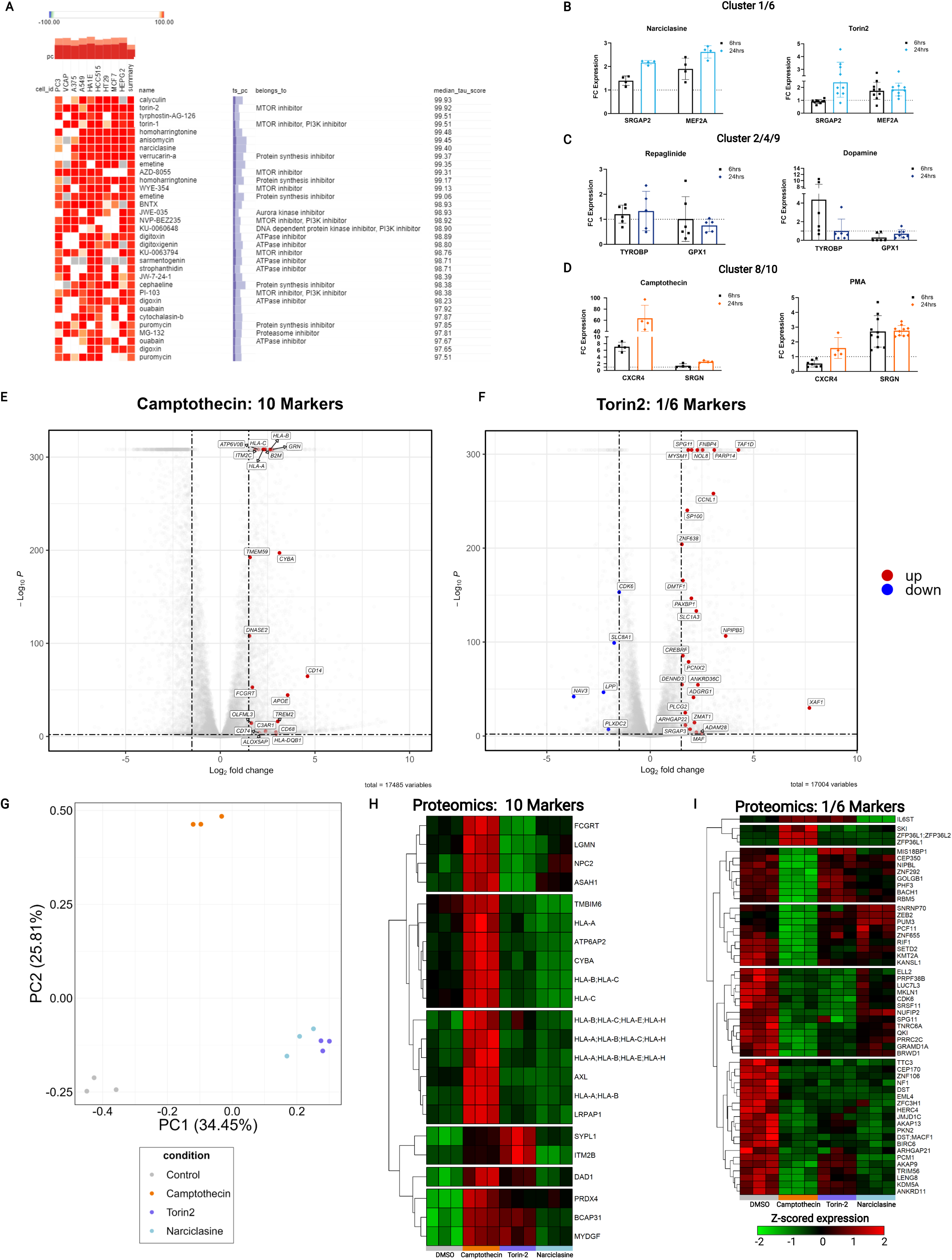
Chemical perturbation recapitulates *in vivo* human microglial subtype signatures *in vitro*. **(A) Representative example of CMAP analysis.** The Connectivity Map (CMAP) was used to identify compounds that might drive transcriptional signatures found in different microglial subsets. The cell_id column identifies the 9 cell lines used in CMAP. Drugs were ranked by the tau score, which quantifies homology between the perturbagen and the query. Scores greater than 90 were considered as candidates for further study. **(B)-(D) qPCR hits by grouping: 1/6, 4/9, or 8/10.** Drugs were tested in the HMC3 microglial model system at 6 and 24h intervals, and two marker genes were assayed by quantitative PCR per cluster group (1/6: *SRGAP2*, *MEF2A*; 4/9; *TYROBP*, *GPX1*, 8/10: *CXCR4*, *SRGN*). CT values were normalized to *HPRT1*. Bars represent fold change expression in relation to DMSO control. **(E) Camptothecin upregulates cluster 10 markers.** Volcano plot showing log fold change (LFC, x), and -log_10_ p-value (y) from bulk RNA-seq generated from HMC3 cells treated with Camptothecin for 24h. Data was analyzed with DESeq2. FDR threshold was set to 0.01 and LFC threshold was set at 1.5. The top 20 cluster 10 genes in the differentially expressed gene list, irrespective of direction, were plotted. **(F) Torin-2 upregulates most cluster 1/6 markers. (G) PCA reveals convergence of Narciclasine and Torin-2 at the proteomic level.** Principal component analysis (PCA) was calculated on log-normalized proteomic data. At the proteomic level, Torin-2 and Narciclasine are similar and divergent from both control and Camptothecin. **(H) Camptothecin upregulates cluster 10 markers at the proteomic level.** Heatmap showing the row-scaled, zero-centered expression values of proteomic data derived from compound-treated HMC3 microglia (24hrs; 3n per treatment). Each column is a single sample, and each row is a single gene. Pairwise differential testing between DMSO control and each of our treated conditions was conducted using Welch’s t-test with the Benjamini-Hochberg correction (FDR alpha < 0.05, LFC **< 1).** (I) Camptothecin downregulates cluster 1/6 markers at the proteomic level. **See also** Figures S7/8 and **Tables S7/8.**

To assess the effects of our selected compounds at broader scale, we repeated our experiment and profiled cells with bulk RNA-seq and proteomics after 24 hours of treatment. On the transcriptional level, Camptothecin induces cluster 8 and 10 genes, such as *HLA-C*, *CXCR4*, and *CYBA* (**Figure 7E**). Interestingly, Camptothecin also downregulates cluster 1/6 genes such as *QKI* and *ATM* (**Figure S7A)**, supporting the transcriptional divergence of clusters 8/10 from 1/6 (**Figure 1C**). As predicted, Torin-2 robustly drives the cluster 1/6 signature (**Figure 7F**). Narciclasine does not appear to upregulate this signature (**Figure S7B**); however, GO annotation of genes differentially upregulated in Narciclasine suggests a strong upregulation of metabolic pathways, such as nitrogen-compound containing metabolism and heterocylic metabolism (**Figure S7C)**, that we previously found to be strongly enriched in clusters 1/6 by GO annotation. Moreover, examining cluster 1/6 genes upregulated in Torin-2 and Narciclasine suggests that the two compounds engage complementary, but separate aspects of the cluster 1/6 signature (**Figure S7D**), with Narciclasine inducing genes such as *MEF2A* and *NUFIP5*, while Torin2 upregulated genes such as *DENND3* and *ATM*. In contrast, at the proteomic level, PCA suggests that Narciclasine and Torin-2 yield similar changes in proteomic profiles relative to both the DMSO control and Camptothecin (**Figure 7G)**, suggesting the engagement of a different proteomic state. Interestingly, neither Narciclasine nor Torin-2 clearly drive cluster 1/6 marker genes at the proteomic level (**Figure 7H**). This may be because RNA-derived markers may be sub-optimal to resolve proteome changes for these microglial subtypes given the known divergence between RNA and protein in microglia^67^ and/or the short time course of our perturbation. On the other hand, Camptothecin does drive strong upregulation of 10 genes such as *HLA*-C and *CYBA* (**Figure 7I**) and downregulation of 1/6 genes such as *QKI* (**Figure 7H**) at the proteomic level.

We have thus identified and validated three tool compounds that polarize a human microglial model system (HMC3 cells) into different targeted states, presenting an approach that can be extended to develop a broader toolkit with which to manipulate microglial differentiation *in vitro* and potentially *in vivo*. Notably, one of our compounds, Camptothecin, drives a robust signal towards cluster 10 that is detectable at both transcriptomic and proteomic levels; a compound with this property could conceptually be useful therapeutically to shift the distribution of human microglia *in vivo* away from clusters 1/6 that are strongly enriched in genes and traits associated with AD, MS, and other diseases (**Figure 4**) and towards the CD74^high^ cluster 10 subset that we have previously reported to be depleted in AD^4^.

## Discussion

Our understanding of human microglial heterogeneity has been transformed over the last 5 years by the efforts of many groups^12, 13^. However, most analyses have adopted a case-control design, where microglia from human donors with a given neuropathological diagnosis are compared to “control” donor microglia. These efforts have been valuable, providing insights into how microglia may contribute to neurodegeneration and neuroinflammation. However, to unify disparate efforts across diseases, there is a need for a common framework, an underlying population structure across different diseases and regions to enable comparisons and translation of results across individual studies. Our goal with this study was to take a first step in this direction; we set out to sample as many microglial subtypes and states as possible and then build a first-generation cross-disease microglial population framework.

In this study, we present an integrated microglial population structure derived from 215,658 live human microglia sampled from a wide array of CNS regions and neurological disease conditions (**Figure 1**). Our analysis explores the interconnection of different microglial subtypes, proposing divergent routes of differentiation with exclusive terminal endpoints as well as possible functional and metabolic shifts associated with these trajectories. We identify families of microglial subsets with enrichment of susceptibility genes for neurodegenerative or neuropsychiatric disease and association with pathological and clinical traits related to AD. To validate our findings, we present a generalizable pipeline combining a joint immunofluorescence & RNAscope staining protocol with automated segmentation to identify transcript abundance in individual microglia, allowing us to confirm the existence of several subtypes *in situ* and to evaluate the association of morphology with gene expression. We also demonstrate the utility of our dataset as a reference by illustrating the interpretation of external data with our cluster model, by assessing its ability to evaluate nucleus-derived data and by evaluating GBM, a disease not captured in our tissue samples. Finally, we demonstrate how a dataset such as this may be used to develop therapies targeting microglial subtypes in the future: we leveraged the CMAP chemical perturbation database and ultimately validated three compounds that each engage one of two examined microglial differentiation programs.

Some of the first studies involving *ex vivo* human microglia have suggested the continuous nature of microglial phenotypes^4, 16^, and indeed, we have found that in our model, shifts in function, metabolism, and association with disease genes or clinicopathological traits fall along continuous axes radiating outwards from a central state that lacks specification beyond general microglia markers. The primary axis of variation in our dataset lies between clusters 1/6 and clusters 2/4/9. Clusters 1/6 are enriched in genetic risk for CNS diseases, associated with clinicopathological traits in AD, and upregulate heterocyclic metabolism. On the other hand, clusters 2/4/9 are associated with oxidative metabolism and present a homeostatic-active phenotype. The second major axis of differentiation separates clusters 8/10 from the other clusters, although they are most distant from 1/6. Both are associated with interferon signaling and T-cell interaction, but cluster 10 shows stronger complement and antigen presentation signatures, while cluster 8 is enriched for steroid and interleukin signaling. These clusters may represent two tracks of microglial activation directed towards adaptive immune interaction, perhaps contingent on the underlying inflammatory milieu. In the final trajectory, the DAM-like trajectory, cluster 11 represents the strongest signal for a DAM2^high^ population yet reported in human data, while cluster 10 shows upregulation of both the DAM1 and DAM2 modules. Interestingly, cluster 9, which is DAM1^high^, also shows a connection to the proliferative cluster, cluster 12. Indeed, it has been previously reported that microglial proliferation is associated with inflammatory stimuli^26^. This suggests that this axis captures a partially overlapping type of activation to our 8/10 axis, one that is suggested in the literature to be heavily associated with neurodegenerative or neuroinflammatory processes. Thus, we have captured in our model three primary axes of variation: (1) a 1/6 *vs.* 2/4/9 metabolic cline, (2) the 8/10 axis of immune response specialization, and (3) the DAM-like axis of activation that terminates in cluster 11. Notably, our *in situ* validation efforts, which avoid the known issues of divergence between RNA and protein in microglia^67^, confirm the existence of these axes by identification of key marker genes. These data also suggest functional differences as we report morphological shifts along some of these axes. For example, CD74^high^ cells (cluster 10) and SPP1^high^ cells (cluster 11) are both less ramified, while CXCR4^+^ (cluster 8) cells exhibit more ramification. This *in situ* experimental and analytic pipeline also offers opportunities to explore spatial localization of RNA in microglia *in situ*, as we can detect RNA transcripts even in distal processes of microglia (**Figure 5**). Further work *in vitro*, *in situ*, and *in vivo* will be required to more comprehensively profile the morphological, metabolic, and functional shifts that occur along these three key axes of differentiation.

Label transfer offers the opportunity to extend our existing structure to external datasets, and our results suggest that both a human iPSC xenograft microglial model system^60^ and an *in vitro* human iPSC microglial model system^62^ recapitulate an impressive amount of the heterogeneity that we observed in tissue-derived live microglia, an encouraging sign for the field. We also recapitulate previously reported findings in these systems, such as an overrepresentation of a plaque and lipid-associated SPP1-high state in the *in vivo* 5XFAD model^61^. Similarly, the analysis of a GBM dataset^58^ generated using a microwell-based scRNA-seq technology^68^ highlights the role of cluster 11 microglia in that context, consistent with prior literature^64^. Finally, despite technical differences, we were able to annotate ROSMAP snRNA-seq data and identified shifts in microglial subsets in association with pathological traits, consistent with our analyses assessing enrichment of disease genes. Ultimately, our dataset and label transfer approach provide a resource for the field that others may use to annotate smaller datasets and enhance the capture of even small numbers of the rarer microglial subtypes that we have identified in our large, unified population model.

While our results from annotating microglial model systems are encouraging, there is still a need for tools that drive and sustain specific microglial phenotypes to enable further study. Our *in silico* analysis prioritized compounds that we validated *in vitro* such as Torin-2, a mTOR inhibitor that improves survival in animal models of GBM^69^, Camptothecin, a topoisomerase inhibitor with neuroprotective effects in murine models of PD^70^, and Narciclasine, a pleiotropic drug that inhibits the NF-κB pathway^71^. Interestingly, mTOR inhibition shifts metabolism in animal models of Leigh Syndrome, exhibiting neuroprotective effects^72^. Similarly, Torin-2 may offer an opportunity to shift modulate microglial metabolism in cases where glycolysis is over-represented. In contrast, Camptothecin is particularly interesting as it robustly enhances its target cluster 10 signature while also suppressing the 1/6 signature at both the transcriptomic and proteomic levels. These effects may be particularly relevant to therapeutic development in AD as clusters 1/6 are enriched for AD susceptibility genes (**Figure 4**) and we have previously reported a depletion of the CD74^high^ cluster 10 in AD^4^. This effort offers a generalizable strategy for identifying compounds that may drive distinct microglial subtypes as a path towards translation and a targeted immunomodulation approach.

Our work has limitations associated with sample accessibility. First, donors of brain tissue are never “healthy”: they are either undergoing autopsy (and are therefore deceased) or a neurosurgical procedure to address a pathologic condition. Thus, we call “control” donors those who have no clinical manifestation of neurological disorder at the time of death and do not fulfill pathological criteria for a disease. Donors in this category are rare as there are few dedicated efforts to collect samples from such participants. Given these constraints, our dataset is not powered to examine shifts in proportion associated with disease states. Similarly, although our analytical pipeline aims to establish a common reference population structure that spans diseases and regions, our efforts to mitigate technical and batch effects likely suppressed some true biological variation among samples. This was a necessary tradeoff to achieve our goal, but it also means that we were unable to capture more subtle heterogeneity that may be present in different diseases or regions. Our dataset may be repurposed by other investigators who wish to further investigate this question in combination with their own data. Moreover, our approach to the analysis of these data began with the standard of single-cell analysis in the field: hard assignment of class labels by way of clustering. However, our data has made it very clear that unlike cell types such as neurons that have discrete, irreversible differentiation endpoints^34^, individual microglia sit along a continuous spectrum with at least three major axes that lead to different states. Finally, our work on recapitulating families of microglial subtypes *in vitro* relies on singular chemical perturbations to drive microglia down different trajectories of differentiation, and we have assessed only relatively short time points. Transcriptional programs that drive the distinct microglial phenotypes that we have observed in this study may require more time and/or combinatorial perturbation to recapitulate more accurately *in vitro* what we observe in live microglia. Indeed, our results with Torin-2 and Narciclasine support the hypothesis that for the most accurate recapitulation, two or three drugs that drive different aspects of a single microglial phenotype may be needed to fully recapitulate the *in vivo* function of a given microglial family. Alternatively, genetic perturbation may be required to drive a stable phenotype to allow for longitudinal study of microglial functions *in vitro*.

Here, we have created a new cross-disease resource investigating heterogeneity of human microglia at the single cell level across a diverse sampling of CNS regions and diseases. Through our validation efforts, we have used it to expand the community’s microglial toolkit with robust approaches to (1) identify microglial subtypes *in situ* and, more broadly, to capture targeted RNA expression from individual microglia in human tissue. Further, (2) we have proposed an approach for label transfer of a microglial reference to multiple data types. Finally, our proof-of-concept demonstration that elements of this microglial heterogeneity can be recapitulated *in vitro* by way of chemical perturbation outlines a path for targeted development of immunomodulatory strategies that leverages our understanding of microglial subtypes implicated in different diseases.

## Acknowledgements

We thank the individuals and their families who donated the brain samples used in this project. The work was supported by the Chan-Zuckerberg Initiative’s Neurodegeneration Challenge Network grant CS-02018-191971. Some of the work also emerged from support from NIH/NIA grants R01 AG070438, U01 AG061356, RF1 AG057473, R01AG048015. Research reported in this publication was supported by the National Institute of General Medical Sciences of the National Institutes of Health under Award Number T32GM007367 and by the National Cancer Institute of the National Institutes of Health under Award Number F30CA261090. The Parkinsonism brain bank at Columbia University is supported by the Parkinson’s Foundation. R.A.H. was supported by grant funding from the Huntington Disease Society of America and Hereditary Disease Foundation and was a Columbia University Irving Medical Center ADRC Research Education Component trainee (P30AG066462). We are grateful to the Banner Sun Health Research Institute Brain and Body Donation Program of Sun City, Arizona for the provision of human brain tissue. The Brain and Body Donation Program has been supported by the National Institute of Neurological Disorders and Stroke (U24NS072026 National Brain and Tissue Resource for Parkinson’s Disease and Related Disorders), the National Institute on Aging (P30AG19610 and P30AG072980 Arizona Alzheimer’s Disease Core Center), the Arizona Department of Health Services (contract 211002, Arizona Alzheimer’s Research Center), the Arizona Biomedical Research Commission (contracts 4001, 0011, 05-901 and 1001 to the Arizona Parkinson’s Disease Consortium) and the Michael J. Fox Foundation for Parkinson’s Research. The Genomics Shared Resource is supported by is supported by NIH/NCI P30CA013696. ROSMAP is supported by P30AG10161, P30AG72975, R01AG15819, R01AG17917. U01AG46152, U01AG61356. ROSMAP resources can be requested at https://www.radc.rush.edu. Research reported in this publication was partially performed in the Columbia Center for Translational Immunology Flow Cytometry Core, supported in part by the Office of the Director, National Institutes of Health under awards S10OD020056.

## Contributions

Conceptualization, J.F.T., P.L.D.; Methodology, J.F.T., M.T., P.A.S., V.M., P.L.D.; Software, J.F.T., M.T.; Validation, J.F.T., M.T., V.H., P.L.D.; Formal Analysis, J.F.T., V.M., P.A.S.; Investigation, J.F.T., M.T., V.H., A.K., M.O., P.L.D.; Resources, B.H., M.F., S.H., T.G.B., J.C., R.K.S., A.F.T., R.A.H, R.N.A, N.S., J.S., D.A.B.; Data Curation, J.F.T., M.O., V.M..; Writing-Original Draft, J.F.T., P.L.D.; Writing-Review & Editing, all authors.; Visualization, J.F.T., V.H.; Supervision, P.L.D.; Project Administration, P.L.D.; Funding Acquisition, P.L.D.

## Materials and Methods

### Source of central nervous system specimens

Details of the acquisition of autopsy samples from Rush University Medical Center/Rush Alzheimer’s Disease Center (RADC)^48, 49^ in Chicago, IL (Dr. Bennett) and Columbia University Medical Center/New York Brain Bank in New York, NY (Drs. Vonsattel and Teich)^73^, as well as surgically resected brain specimens from Brigham and Women’s Hospital in Boston, MA (Drs. Sarkis, Cosgrove, Helgager, Golden, and Pennell) were detailed in our prior publication^4^. In addition, samples were obtained from donation programs at Massachusetts General Hospital, Boston, MA (Drs. Bradley T. Hyman and Matthew Frosch), Banner Sun Health Research Institute, Sun City, AZ (Dr. Thomas G Beach) and Rocky Mountain MS Center, Denver, CO (Dr. John Corboy). All brain specimens were obtained through informed consent and/or brain donation program at the respective organizations. All procedures and research protocols were approved by the corresponding ethical committees of our collaborator’s institutions as well as the Institutional Review Board (IRB) of Columbia University Medical Center (protocol AAAR4962). For a detailed description of the brain regions sampled, clinical diagnosis, age and sex of the donors see **Table S1**.

### Shipping of brain specimens

After weighing, the tissue was placed in ice-cold transportation medium (Hibernate-A medium (Gibco, A1247501) containing 1% B27 serum-free supplement (Gibco, 17504044) and 1% GlutaMax (Gibco, 35050061)) and shipped overnight at 4°C with priority shipping.

### Microglia isolation, cell hashing and sorting

The isolation of microglia was performed according to our published protocol^8^, with minor modifications. In case of the cortical autopsy samples (BA9/46, BA4, BA17/18/19), the cortex (grey matter and the underlying white matter (subcortical white matter) were dissected under a stereomicroscope. The subcortical white matter samples were not used in this study. The epilepsy surgery samples of temporal lobe (BA20/21) were processed without dissection as in this case the cortical white and grey matter was not always distinguishable due to the surgical procedure. The substantia nigra (SN) and the thalamus (TH) were dissected by separation from the surrounding white matter tracts. The hippocampus samples (H) contained the dentate gyrus, CA4/CA3/CA2 and CA1 regions, both white and grey matter. The spinal cord sample (SC) was sampled at the level of lumbar section and included both white and grey matter. The anterior watershed area (AWS) deep white matter did not need any further dissection. All steps of the protocol were performed on ice. The dissected tissue was placed in HBSS (Lonza, 10-508F) and weighed. Subsequently the tissue was homogenized in a 15 ml glass tissue grinder - 0.5 g at a time. The resulting homogenate was filtered through a 70 um filter and spun down at 300rcf for 10 minutes. The pellet was resuspended in 2 ml staining buffer (RPMI (Fisher, 72400120) containing 1% B27) per 0.5 g of initial tissue and incubated with anti-myelin magnetic beads (Miltenyi, 130-096-733) for 15 minutes according to the manufacturer’s specification. The homogenate was than washed once with staining buffer and the myelin was depleted using Miltenyi large separation columns (Miltenyi, 130-042-202). The cell suspension was spun down and was then incubated with anti-CD11b AlexaFluor488 (BioLegend, 301318) and anti-CD45 AlexaFluor647 (BioLegend, 304018) antibodies as well as 7AAD (BD Pharmingen, 559925) and cell hashing antibodies (for catalogue numbers of cell hashing antibodies see Table S1) for 20 minutes on ice. Subsequently the cell suspension was washed twice with staining buffer, filtered through a 70 µm filter and the CD11b+/CD45+/7AAD- cells or CD45+/7AAD- cells were sorted on a BD FACS Aria II or BD Influx cell sorter. Cells from each brain region were sorted in a separate A1 well of a 96 well PCR plate (Eppendorf, 951020401) containing 100 µl of PBS buffer with 0.3% BSA. Following sorting cell from different brain regions were combined and immediately submitted to single cell capture, reverse transcription and library construction on the 10x Chromium platform. All sorting was performed using a 100 µm nozzle. The sorting times varied according to the quality of the sample but was on average between 10 and 20 minutes per sample. The sorting speed was kept between 8000 - 10,000 events per second.

### 10x Genomics Chromium single cell 3’ library construction

Cell capture, amplification and library construction on the 10x Genomics Chromium platform was performed according to the manufacturer’s publicly available protocol. Briefly, viability was assessed by trypan blue exclusion assay, and cell density was adjusted to 175 cells per μl. 7,000 cells were then loaded onto a single channel of a 10x Chromium chip for each sample. The 10x Genomics Chromium technology enables 3’ digital gene expression profiling of thousands of cells from a single sample by separately indexing each cell’s transcriptome. First, thousands of cells are partitioned into nanoliter-scale Gel Bead-In-EMulsions (GEMs). Within one GEM all generated cDNA share a common 10x barcode. Libraries were generated and sequenced from the cDNA, and the 10x barcodes were used to associate individual reads back to the individual partitions. To achieve single cell resolution, the cells were delivered at a limiting dilution. Upon dissolution of the Single Cell 3’ Gel Bead in a GEM, primers containing (i) an Illumina R1 sequence (read 1 sequencing primer), (ii) a 16 nucleotide 10x Barcode, (iii) a 10 nucleotide Unique Molecular Identifier (UMI), and (iv) a poly-dT primer sequence were released and mixed with cell lysate and Master Mix. Incubation of the GEMs then produced barcoded, full-length cDNA from poly-adenylated mRNA. After incubation, the GEMs were broken and the pooled fractions were recovered. Full-length, barcoded cDNA was then amplified by PCR to generate sufficient mass for library construction. Enzymatic fragmentation and size selection were used to optimize the cDNA amplicon size prior to library construction. R1 (read 1 primer sequence) were added to the molecules during GEM incubation. P5, P7, a sample index, and R2 (read 2 primer sequence) were added during library construction via end repair, A-tailing, adaptor ligation, and PCR. The final libraries contained the P5 and P7 primers used in Illumina bridge amplification. The described protocol produced Illumina-ready sequencing libraries. A Single Cell 3’ Library comprises standard Illumina paired-end constructs which begin and end with P5 and P7. The Single Cell 3’ 16 bp 10x Barcode and 10 bp UMI are encoded in Read 1, while Read 2 is used to sequence the cDNA fragment. Sample index sequences were incorporated as the i7 index read. Read 1 and Read 2 are standard Illumina sequencing primer sites used in paired-end sequencing. Sequencing the library produced a standard Illumina BCL data output folder. The BCL data includes the paired-end Read 1 (containing the 16 bp 10x Barcode and 10 bp UMI) and Read 2 and the sample index in the i7 index read.

### Batch structure and sequencing

Tissue specimens were processed upon receipt. The different brain regions from the same donor were processed and hashed parallel and loaded in a single well of a 10x Chromium 3’ chip as described above. Accordingly, each sample (containing multiple brain regions from the same donor) constitutes one batch for all three procedures (microglia isolation, cell capture and library construction). All sequencing was performed on an Illumina HiSeq4000 or a NovaSeq6000 machine. For specifics on the sequencing machines and QC metrics regarding the generated reads see. Table S1.

### Single-cell RNA-seq data processing, alignment, and hashtag deconvolution

The majority of our downstream analysis was conducted using the R programming language (v4.0.5 for harmonization and clustering, v4.1.0 for annotation and downstream visualization)^74^ and the RStudio^75^ integrated development environment. CellRanger V3.1.0 with default parameters was used to demultiplex and align our barcoded reads with the Ensembl transcriptome annotation (downloaded March 2019, GRCh38.91). A recent report^76^ suggested that filtering cells with greater than 10% mitochondrial reads is the preferred baseline for human tissue, and that for brain tissue a higher threshold may even be optimal. Thus, a mitochondrial percentage that was the higher of either 10% of reads or the 2 absolute deviations above the median for mitochondrial reads within the sample was chosen as a threshold. Cells below this threshold with between 500 and 10,000 UMIs were retained for downstream analysis. All ribosomal genes, mitochondrial genes, and pseudogenes were removed, as they interfered with the downstream differential gene expression. For samples where we used cell hashing to combine regions or subjects in a single sequencing run, droplets were demultiplexed using the following workflow. For each HTO, a mixture model with two components was fitted to the HTO counts using an EM algorithm. The component with the smaller mean (negative component) represents droplets that were not tagged with the HTO, whereas the component with the larger mean (positive components) represents droplets that were tagged. We then assign each droplet to either the negative or positive component based on its posterior probability. Droplets that were assigned to the negative component for all HTOs as well as multiplets were discarded. Singlets with high uncertainty, i.e. without confident assignment to either the negative or positive component, were discarded as well, leaving only high certainty singlets for downstream analysis. The method is implemented in the R package demuxmix which available on github: https://github.com/cu-ctcn/demuxmix. Some of our hashtag data had lower overall counts, and thus, the demuxmix model was unable to effectively segregate distributions for some hashtags in several samples. These samples were identified as having high percentages of negative/uncertain cells with demuxmix. In these cases, to try and recover cells for further analysis, the problematic hashtags were reclassified using one of two different algorithms, a demixing algorithm developed for MULTI-seq^77^ or HTOdemux from Seurat v3.2.0^23^. Hashtag classifications were merged, and doublet/negative/uncertain cell removal proceeded as before.

### Batch correction

Striking differences were observed in the distributions of UMI counts between 10X v2 and v3 chemistry. As this was driving differential clustering, count matrices from v3 samples were downsampled by 50% using the DropletUtils^78^ package in R to achieve comparable UMI distributions across the two technologies (Table S1, Figure S2D). Next, after testing a series of recently published batch correction tools, SCTransform^79^ combined with mNN^80^ was chosen to mitigate batch effect in our dataset. A range of numbers of differentially expressed genes (3000-6000) and components (20-40) were tested, and 4500 differentially expressed genes and 40 components were used for downstream analysis. Using these parameters, the full pipeline is as follows: SCTransform, which normalizes for minor differences in sequencing depth, was performed on each individual batch, then corrected counts were log-normalized with the *NormalizeData* function in the Seurat package. Subsequently, processed datasets were merged on the corrected count matrix using the fastmNN algorithm accessed through the *RunFastMNN* function in the SeuratWrappers package. Library preparation batch is confounded with sample, and diseases are confounded with technical variables in our dataset, due to the necessity to process all tissue immediately upon receipt and the irregular schedule by which samples are received. As mNN is a harsher integration approach when compared to other commonly used tools, our pipeline is likely to have removed relevant biological signal in integrating the datasets. However, our priority was to obtain a robust cluster structure across the diverse brain regions and diseases found in our dataset while avoiding the issue of spurious signal from batch effect driving separate clustering, which motivated the approach we have described here.

### Clustering

The graph-based clustering approach implemented in Seurat V3^23^ was used to cluster our cells. In brief, a KNN graph based on Euclidean distance in our corrected mNN space was calculated and used to derive refined edge weights based on Jaccard similarity. The Louvain algorithm was then applied to iteratively delineate a population structure on our dataset. This was implemented with the *FindNeighbors* and *FindClusters* functions in Seurat. A UMAP projection of our dataset was computed with the *RunUMAP* function for visualization (Figure S1A). Contaminating red blood cells from our dataset were removed using classical markers (*HBB/HBA*) and microglia were subclustered using an identical integration and clustering pipeline (Figure 1A). Any microglial subsets with fewer than 100 cells were discarded. Basic quality metrics are shown in Figure S2A-F and reported in Table S1.

### Validation of cluster stability

To evaluate cluster stability, a post-hoc pairwise machine learning approach was used to evaluate the similarity between clusters. The top 10,000 variable genes in the dataset were identified by applying the *FindVariableFeatures* function from Seurat on the log-normalized RNA expression data from our dataset. Using these features, keras^81^ was used to train a multilayer perceptron classifier to distinguish each pair of clusters. After basic hyperparameter optimization, the following parameters were chosen for the model: the rmsprop optimizer, a 1-layer structure with 500 nodes in the hidden layer, the tanh activation function for the dense layer, and a sigmoid activation function for the output layer. 10 epochs and a batch size of 100 were used for training. As our only concern was the raw accuracy of classification, mean squared error was used as the loss function. For each pair of clusters, the data was split into 5 folds, then the classifier was trained on 80% of the dataset and the remaining 20% was classified. This process was repeated 5 times to classify every cell in our dataset once. This entire pipeline was then repeated 50 times for each pair of clusters. Cells that were ambiguously classified (<40 times out of 50) to the same cluster were designated intermediate cells. A threshold of <20% overlap between clusters was chosen as the threshold for merging clusters; in our model, no clusters met this parameter and as such, the original 12-cluster model was retained. The constellation diagram shown in Figure 2A depicts the results of this analysis: edges between clusters represent the percentage of total cells classified as intermediate, and the area of nodes is scaled to correspond to the overall size of the corresponding cluster. As orthogonal validation of our microglial clustering parameters, a resampling clustering approach was also used to assess cluster robustness. Over 100 iterations, 75% of the cells from our dataset were randomly sampled and our clustering pipeline was run with identical parameters. This was also done in a pairwise fashion to examine fluidity between individual pairs of clusters. For each pair of clusters, the frequency with which cells assigned to one cluster in our original clustering were re-clustered into the cluster that contained fewer cells with the same original classification was recorded. Clusters remain generally stable, with most of the overlap being found between adjacent, closely related clusters or the intermediate clusters in our dataset. The results are shown in Figure S2G, visualized using the corrplot package^82^. Each of our microglial subtypes was also subclustered, but no stable, distinct subclusters were identified.

### Identification of cluster-defining gene sets

To identify cluster-defining gene sets, the *FindMarkers* function in Seurat was used to implement a pairwise testing approach. We prioritized differentially expressed genes that could best delineate a given cluster from each other cluster in our dataset. To do so, MAST^83^ was applied to normalized count data from the “RNA” assay of the Seurat object to find differentially expressed genes between every combination of pairs of clusters. Within each cluster, all the differentially expressed genes that were identified with this approach were filtered to only include those that were only found to be differentially expressed in one direction (either up or down). Any genes that were found to be upregulated in comparison to some clusters but downregulated in comparisons to other clusters or vice versa were removed from our downstream analysis. Furthermore, to ensure that the specific cluster-defining genes were prioritized, upregulated genes were ranked by the number of comparisons in which they were upregulated, and only those upregulated in 3 or more comparisons were used for downstream analyses. An identical process was applied for downregulated genes. Full marker gene lists are reported in Table S2.

### Enrichment of disease-associated microglia signature gene sets

The disease-associated microglia (DAM) signature gene sets for “DAM1”, “DAM2”, and “homeostatic microglia” described in Keren-Shaul et al.^6^ were examined separately in our analysis. These sets consist of two sets of genes upregulated in the DAM trajectory, as well as homeostatic microglial genes known to be downregulated in DAM microglia. The overlap of “DAM1” and “DAM2” gene sets with upregulated cluster-defining genes, and overlap of the “homeostatic microglia” gene set with downregulated cluster-defining genes (see Identification of cluster-defining gene sets) was examined using a hypergeometric test with an FDR-corrected threshold for significance at *q* = 0.01^84^. The results of this analysis were visualized in a heatmap where the color intensity corresponds to the -log_10_ p-value of the FDR q-value for enrichment of DAM1/DAM2 genes in upregulated genes or homeostatic genes in downregulated genes from our clusters (Figure 1E).

### Monocle3 pseudotime analysis

As an orthogonal method of evaluating the continuity of different microglial states in our cluster structure, the monocle3 algorithm was used to build a pseudotime trajectory across our dataset as shown in Figure S4E. Using the Seurat interface to monocle3 found in SeuratWrappers, the Seurat object was converted into a monocle data object, and a pseudotime trajectory was derived using the *learn_graph* function, retaining the clustering assignments from our original clustering pipeline (see Clustering).To establish an originating point, the pseudotime root was placed on the border of clusters 2 and 3, as these cells had the strongest gene expression signature for classic homeostatic microglia and few differentiating genes, suggesting that they could form the basal state for microglia from which they would differentiate into other states. Interestingly, this state was best captured by choosing cells with maximal *AVP* expression, a marker of hematopoietic stem cells^85^ that is frequently used to mark the root cells in hematopoietic pseudotime tracing.

### Functional annotation of microglial clusters

To perform functional annotation of microglial clusters, up- or down-regulated gene lists for each cluster were defined as genes upregulated in 3 or more comparisons or downregulated in 2 or more comparisons. Annotation of these gene lists was performed with several resources: Gene Ontology by way of TopGO^37^ as well as Reactome^39^ pathway analysis with clusterProfiler^41^. For GO analysis, we conducted analysis with biological process annotation. For all functional analysis, the Benjamini-Hochberg correction^86^ was used to correct p-values for multiple testing. Corrected p-values below a threshold of 0.01 were chosen as significant for both GO and Reactome results. GO results were aggregated and summarized by use of the rrvgo^38^ package. Aggregated results of pathways are shown in Figure 2 and Figure S4. For Figure 2A, terms were filtered to include only those terms that were simultaneously upregulated in both clusters 4 and 9 and downregulated in both clusters 1 and 6, or vice versa to best highlight differences between these families.

### Examining enrichment of multiple sclerosis susceptibility genes

The enrichment of MS susceptibility genes was evaluated separately from other diseases due to the availability of a recent publication by the International Multiple Sclerosis Genetics Consortium extensively mapping genomic risk loci in MS^45^. A hypergeometric test was used to evaluate the enrichment of 551 putative MS susceptibility genes identified as targets of MS variants in genes upregulated in our clusters (see Identification of cluster-defining gene sets). The FDR-corrected threshold for significance was set at *q* = 0.01.

### Examining enrichment of disease genes from the GWAS Catalog

To confirm the results of our MS analysis and to examine enrichment of genetic risk from other neurodegenerative or neuroinflammatory diseases, we were interested in using a more comprehensive source of disease-gene associations. Thus, the GWAS Catalog^46, 87, 88^, a curated database that focused on SNP-trait association, was used for further analysis. This dataset contains select studies that include a primary GWAS analysis (as per the GWAS catalog website: “array-based genotyping and analysis of 1000,000 pre-QC SNPs selected to tag variation across the genome and without regard to gene content”) or an imputation analysis with sufficient genome-wide coverage to meet the definition of a GWAS catalog mentioned previously. This catalog is updated on a weekly basis by curators, and eligible studies are generally added within 1-2 months of publication. The 2021-08-16 data release was used for this study. For our analysis, we chose to focus on specific disease entities where microglia are proposed to play relevant roles. To narrow our scope, we GWAS catalog entries were filtered by a specific disease name. For example, to examine Alzheimer’s Disease associations, all records containing “Alzheimer” in the “DISEASE_TRAIT” column were retained. Similarly, for stroke and cerebrovascular disorders, we filtered for all records containing the keywords “stroke”, “brain ischemia”, “cerebral ischemia”, “cerebral artery”, and “cerebrovascular”. We carried out a similar approach for all other diseases we investigated in this analysis. After obtaining sets of disease-gene associations for each disease entity of interest, we applied a similar testing approach to that described in our MS disease gene analysis (see Examining enrichment of multiple sclerosis susceptibility genes).

### Examining association of ROSMAP traits with clusters

The ROSMAP RNA-seq cohort used in our analysis contains a total of 1092 samples, with a total of 18,629 genes captured across all samples. Using the DEseq2^89^ R package, the *DESeq* function was used to perform differential expression analysis in association with one of 12 traits. The model for differential testing was:

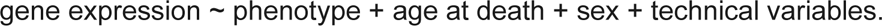

Technical variables included: Batch, LOG_ESTIMATED_LIBRARY_SIZE, LOG_PF_READS_ALIGNED, PCT_CODING_BASES, PCT_INTERGENIC_BASES, PCT_PF_READS_ALIGNED, PCT_RIBOSOMAL_BASES, PCT_UTR_BASES, PERCENT_DUPLICATION, MEDIAN_3PRIME_BIAS, MEDIAN_5PRIME_TO_3PRIME_BIAS, MEDIAN_CV_COVERAGE, pmi, and study. For slope of cognitive decline, additional adjustment was performed for years of education in the model. Lists of positively and negatively associated genes were derived with an FDR-adjusted p-value with alpha = 0.01. Enrichment of positively and negatively associated genes for each trait in the up- and down-regulated genes for each cluster respectively was evaluated with a similar testing approach and threshold to our disease annotation analyses (see Examining enrichment of disease genes from DisGeNET). Detailed descriptions of the ROSMAP traits used in this analysis can be found at https://www.radc.rush.edu/docs/var/varIndex.htm, and they have been described in detail elsewhere^8, 48, 49, 90, 91^. Results of this analysis are reported in Table S4.

### In situ confirmation of microglia subset identity and abundance

As data does not always correlate between the transcriptomic and protein levels, a phenomenon that has been noted to be pronounced specifically in activated microglia^67^, Advanced Cell Diagnostic’s RNAscope was used to confirm our scRNA-seq results. A cohort of samples from the New York Brain Bank (details on donor cohort can be found in Table S5) consisting of prefrontal cortex (BA9) tissue sections from 16 donors was chosen for validation efforts. After extensive optimization of a co-detection workflow to merge immunofluorescence (IF) and RNAscope, our final pipeline is described below. Initial optimization was performed with positive and negative RNAscope 4-plex controls (ACD, Positive: 321831, Negative: 321801), and once an optimal pipeline was identified, it was run with probes of interest.

All reagents from the RNAscope Multiplex Fluorescent Reagent Kit v2 (ACD, 323100) were prepared for use in accordance with the manufacturer’s instructions. All wash buffers were prepared immediately prior to performing experiments.

6 µM tissue sections were deparaffinized with CitriSolv Clearing Agent (Decon Laboratories, Inc., 1601) for 20 min at room temperature (RT). This was followed by an ethanol series (100%, 100%, 70%) (100%: Fisher Scientific, BP2818, 70%: Fisher Scientific, BP8203) for 30 sec per bath with agitation and re-hydration in distilled water for 1 minute at RT. Subsequently, 4-6 drops of Hydrogen Peroxide were used to cover the tissue section, and slides were incubated for 10 min at RT. Hydrogen peroxide (ACD, 322381) solution was removed by tapping on absorbent paper and slides were washed with distilled water twice. Antigen retrieval was performed with pH 6.0 citrate (Sigma-Aldrich, C9999) and heating with a microwave for 25 min at 400 watts. Slides were then placed in tap water for 5 minutes, then moved to 100% ethanol for 1 minute. Slides were allowed to dry fully at RT, then hydrophobic barriers were drawn around the tissue section with Super Pap Pen Liquid Blocker (Newcomer Supply, 6505). Slides were then blocked for 30 minutes at RT with RNAscope Co-Detection Antibody Diluent (ACD, 323160). Diluent was removed by tapping on absorbent paper, and slides were treated with primary antibody diluted in RNAscope Co-Detection Antibody Diluent for 2 hrs at RT. Slides were washed three-times with PBS (Corning, 46-013-CM) containing 0.1% Tween-20 (Sigma-Aldrich, P9416) (PBS-T), then submerged in fresh 10% Neutral Buffered Formalin (Sigma-Aldrich, HT5011) for 1 hr at RT. Slides were washed with PBS-T 3 times, then 4 drops of RNAscope protease plus (ACD, 322381) were added to the slide and spread to fully cover the tissue. After incubation for 40 min at 40°C in the RNAscope HybEz II oven (ACD, 321710), slides were washed once with distilled water. 125 µL of pre-mixed RNAscope probe mix was then added to each slide, and then slides were incubated for 2 hr at 40°C. Slides were removed from the oven and washed twice with RNAscope wash buffer (ACD, 310091). Slides were then covered with 5X SSC (Sigma-Aldrich, S6639-1L) buffer and left overnight until the morning, when the protocol was resumed.

On the second day of the protocol, the slides were washed twice with RNAscope wash buffer, then 4 drops of RNAscope AMP1 (ACD, 323101) were added to each slide. After 30 min of incubation at 40°C, slides were washed twice with RNAscope buffer, then 4 drops of AMP2 (ACD, 323102) were added per slide. After 30 more min of incubation at 40°C, slides were washed twice with RNAscope buffer, then 4 drops of AMP3 (ACD, 323103) were added per slide. After a final 15 minutes of incubation, slides were washed twice with RNAscope buffer.

Next, 4 drops of HRP-C1 (ACD, 323104) were added per slide. After 15 min of incubation at 40°C, slides were washed twice with RNAscope buffer, then 150 µL of Opal 570 (Akoya, FP1488001KT) dye diluted in RNAscope TSA diluent (ACD, 322809) were added per slide. Slides were incubated for 30 minutes, then washed twice with RNAscope wash buffer. Finally, 4 drops of HRP blocker (ACD, 323107) were added, followed by a 15-minute incubation period and two washes with RNAscope buffer. This HRP-TSA-block process was repeated identically with either HRP-C2 (ACD, 323105) or HRP-C3 (ACD, 323106) depending on the channel of the original probes, and Opal 690 (ACD, FP1497001KT) dye diluted in TSA diluent. Finally, this HRP-TSA-block process was repeated once more with HRP-C4 (ACD, 323121), and the following modifications: TSA-DIG (Akoya, FP1501001KT) diluted in RNAscope TSA diluent for the second step and a 30 min incubation at RT instead of 40°C, and swapping the second wash after HRP blocking from RNAscope wash buffer to PBS-T.

After all HRP steps were completed, counterstaining was performed with 200 µL of secondary antibody diluted in RNAscope Co-Detection Antibody Diluent. After incubation for 30 min at RT, slides were washed three times with PBS-T. Slides were then incubated with 150µL of Opal Polaris 780 dye (Akoya, FP1501001KT) diluted in Antibody Diluent/Block (Akoya, ARD1001EA) for 30min at RT. After three washes with PBS-T, slides were incubated with 200µL of 1X Trueblack (Biotium, 23007) for 2 min at RT to quench lipofuscin autofluorescence. Slides were washed three times with PBS and counterstained with 4 drops of DAPI (ACD, 323108) per slide incubated at RT for 30 seconds. After removing DAPI by tapping slides on absorbent paper, the hydrophobic barrier was removed, and the slides were mounted with 1 drop of Prolong Gold (Thermo Fisher Scientific, P36934) and coverslips (Fisher Scientific, 12545F). Bubbles were removed from the mounting medium using gentle pressure from a pipette tip. Slides were dried for 30 min at RT in the dark, then transferred to 4°C for imaging the following day.

The primary antibody used in staining was goat anti-human Iba1 (Wako, 01127991; dilution of 1:50), the secondary antibody used in staining was donkey anti-goat IgG (H+L) highly cross-adsorbed secondary antibody conjugated to Alexa Fluor Plus 488 (ThermoFisher Scientific, A11055;1:500). The RNAscope probes used in our experiments were: *CD74* (ACD, 477521), *CXCR4* (ACD, 310511-C2), *GPX1* (ACD, 492881), and *SPP1* (ACD, 420101-C4). Two additional probes were used but provided insufficient signal for downstream analysis: *MEF2A* (ACD, 452891-C3) and *CX3CR1* (ACD, 411251-C3).

Fields of view were captured using the 40x objective of a Nikon Eclipse Ni-E immunofluorescent microscope. For each donor, 15 images were obtained from the gray matter with same exposure time, then loaded into CellProfiler software where automated segmentation and downstream analysis was performed as described below. Representative images can be found in Figure 5B/F and Figure S5A/B.

### Automated image analysis using CellProfiler

To automatically segment images and localize transcripts within microglia, we developed an extensive pipeline in CellProfiler v4.2.1. First, the *IdentifyPrimaryObjects* module was used to segment based on DAPI, the *EnhanceOrSuppressFeatures* module was used to enhance the Iba1 signal (using “Neurites” as the feature type) using the “Line structures” filter, and segmented Iba1 signal was identified using *IdentifyPrimaryObjects*. After another round of enhancement of Iba1 signal by applying the “Tubeness” filter (again using “Neurites” as the feature type), the *RelateObjects* module was used to relate segmented DAPI and Iba1 objects using the DAPI as the parent objects and the Iba1 (after enhancement) as the child objects. Next, morphological parameters of each joint-segmentation defined cell were measured with the *MeasureObjectSizeShape* module. Then, for each of the channels with RNAscope imaging data, the following set of steps were run: enhancement of the signal for the given channel with *EnhanceOrSuppressFeatures* specifying “Speckles” as the feature type, identification of RNA puncta using *IdentifyPrimaryObjects*, setting a mask for the channel with *MaskObjects*, then relating identified RNA puncta to the parent cells with *RelateObjects*. Subsequently, the intensity in each of the channels for all RNA that were associated with a parent microglial cell was measured with the *MeasureObjectIntensity* module, and all data was exported with the *ExportToSpreadsheet* module. Our full pipeline is available in Table S5.

### Analyzing CellProfiler-processed RNAscope data

Downstream of CellProfiler processing, all RNAscope puncta in any channel were filtered to exclude those found outside of microglia identified by the joint segmentation pipeline. Using the microglia-localized puncta, an overall area-adjusted score was computed for each channel (cy3, cy5, or cy7) by dividing the summed intensity of puncta detected within a given microglial cell by the computed area for each microglial cell segmented by CellProfiler. The distribution of area-adjusted intensity for the different images in our cohort was evaluated, showing that most of our sections had similar distributions of cells, with rare outliers that had abnormally high signal in all channels. These outliers were excluded from further analysis. For three of our markers, *CD74*, *GPX1*, and *SPP1*, expression levels for each of these markers were thresholded into three bins: low, medium, and high, as we found substantial detection of these markers across all our tissue. In contrast, *CXCR4* was detected only in a small fraction of cells at similar levels, so cells were segmented as either *CXCR4* positive or negative.

Thresholds were determined based on the distributions of the RNAscope data, as well as the levels of expression for different subtypes in our scRNA-sequencing dataset. For example, in our scRNA-seq data, the “high” subtype for *CD74*, cluster 10, has a median fold change of 2.899 for *CD74* compared to other subsets. Thus, the threshold for *CD74* in the RNAscope data was set as 2.899 times the median of *CD74* expression. Similarly, to derive the low threshold for *CD74*, the fold change in *CD74* expression was compared between the set of families with low *CD74* expression, which included the closely related clusters 1, 5, 6, and 7, as well as the proliferative cluster 12, and all other clusters where the difference in expression of *CD74* was found to be significant by our pairwise testing approach (See Identification of cluster-defining gene sets). In this case, the median fold change in expression for our low classes versus all other clusters was 0.394. As such, the *CD74*-low class threshold was set as 0.394 times the median of *CD74* expression. This process was repeated for *SPP1* and *GPX1*; for example, the *GPX1*-high classes were clusters 2, 4 and 9, while the low classes were 1, 6, 7, and 12, and median fold change of these two groups of clusters versus other clusters was used to determine the high and low threshold respectively. A small fraction of cells with abnormally high signal that no longer appeared punctate in form, but rather diffuse and sometimes extending beyond the boundary of the cells was identified. Although these could represent real cells, these might also represent cells with high levels of background in our specific channels. Thus, for *CD74* and *SPP1*, the two markers where these types of cells were observed, cells that were 1.5 IQR above the 75^th^ percentile for expression for all channels were excluded. This excluded a small number of cells (49 out of a total of 7364 cells for panel 1 and 13 out of a total of 3710 cells for panel 2).

To provide the most accurate comparison of numbers of cells between RNAscope and scRNA-seq, cells in scRNA-seq coming from either AD diagnoses (EOAD, LOAD), PD diagnoses (PD-DLBD, PSP), or our control sample, were chosen for comparison, as these were the diagnoses represented in the samples that we obtained for our *in situ* analysis. Proportions of cells per binned class (i.e. *CD74^low^*, *CD74^medium^*, *CD74^high^*) were then compared between the two datasets.

For analyses leveraging various features output by our CellProfiler pipeline, including the “Compactness” and “Eccentricity” features, the output of CellProfiler for each of these features was used. For others, such as “median distance”, the median distance of puncta for a given channel (e.g. cy3, cy5, or cy7) was manually computed from the centroid of a single segmented microglial cells. In all cases involving median distance of puncta from cellular centroids, we excluded all cells in the “low” class for all channels in question to only include cells with real data. Significance of differences in morphological features between expression classes, was tested with Welch’s t-test with the Holm-Bonferroni correction^92, 93^, setting a significance threshold for adjusted p-value of 0.05.

### Training machine learning models for label transfer to other single-cell microglial datasets

In initial evaluation of our query datasets, substantial batch effect was evident. As this was likely to confound our downstream label transfer workflow, a version of our mNN integration pipeline was adapted for upstream removal of batch effect. To do so, our reference data was concatenated with the query data in a single Seurat object and the unique differentially expressed genes from our pairwise differential expression testing (Identification of cluster-defining gene sets) were used for cross-batch merging with the fastmNN algorithm. For all these analyses, we used 40 components. The normalized “mnn.reconstructed” assay, which represents per-gene corrected log-expression values, was used for downstream analysis.

After testing a number of different models in our label transfer pipeline, a combinatorial workflow leveraging two distinct models for different clusters showed the best accuracy: a set of pairwise support vector machine (SVM) classifiers using consensus voting to assign labels for the smaller clusters (8-12), and a flat XGBoost^63^ (XGB) classifier to assign labels for the larger clusters (1-7) with higher transcriptional homology. This set of models was chosen because the SVM achieved highest accuracy in initial testing with smaller classes, but lower-than-average accuracy on larger classes, whereas the XGB results followed the exact opposite trend. Predictions from these two models were thus integrated to achieve higher predictive accuracy. The overall workflow for both methods was similar: as a few our classes are transcriptionally similar, similar classes are condensed (clusters 1/6/7 and clusters 2/4), then a subset of the cells in our dataset are selected for training. Next, the differentially expressed genes from our pairwise differential expression testing (Identification of cluster-defining gene sets) were selected as the features for training and PCA was performed on the resulting subset of the data.

For the SVM, the training subset was 0.2 for classes 1-9, and 0.5 for classes 10-12. A separate classifier was trained for each unique pair of clusters (i.e. a classifier to compare clusters 1/6/7 and 2/4, 1/6/7 and 3….1/6/7 and 12, then 2/4 and 3, 2/4 and 5….2/4 and 12) using only the genes found to be differentially expressed (both up and down) between that specific pair of clusters. Data classes were then rebalanced using combined over/under resampling to reduce class imbalance for smaller classes. Caret^94^ was used to perform PCA and hyperparameter optimization of a SVM model using a radial kernel and 10-fold cross-validation repeated three times. PCA was conducted independently during each fold. Conversely, for XGB, the training subset was 0.33, and the model trained only on cells from groupings 1/6/7, 2/4, 3, and 5. Similarly, PCA was performed upstream on the subset of scaled data consisting of all genes found to be differentially expressed between any clusters. Hyperparameter optimization with 5-fold validation was performed in a stepwise fashion: tree number was first optimized, then tree-specific parameters were tuned with a restrictive grid search, then regularization parameters were tuned with a restrictive grid search, then final optimization was conducted with grid search in a narrow range around prior optimal parameters.

To construct a validation subset, a subset of 50% of the dataset was sampled exclusively from cells not used for training of either the SVM or XGB models. The same scaling and subsetting operations described above were applied to this data. Optimized SVM and XGB models were used to classify the data. For SVM models, final classifications were obtained with hard consensus voting, as the class with the majority of votes was chosen as the final class of the SVM voting ensemble. Similarly, for XGB, which outputs a probability for each class summing to 1 across all classes, the highest probability was used to choose the assigned label. However, the class probabilities for XGB also provided the opportunity to evaluate the confidence of the classifier and drop lower-confidence assignments. As such, cells were only retained for SVM classifications in classes 10-12 or for XGB classifications in classes 1-7 that had higher than 50% classification probability for the assigned probability. Final classifications were merged across datasets, and accuracy was evaluated by examining sensitivity, specificity, and congruence of marker gene expression patterns of cells assigned to each class with marker gene expression patterns seen in our original data. Identical procedures were performed for query datasets.

This approach demonstrates high sensitivity and specificity on test data, with joint accuracy averaging 85% across models trained for different query datasets. Notably, uniformly high specificity is observed, even for clusters with lower sensitivity, such as clusters 3 and 5. These two clusters are also associated with lower confidence scores from our XGB model, an expected result given the transcriptionally intermediate nature of these clusters. Thus, the model’s greatest difficulties with classification come in cases where the true classification boundary is not well defined, which provides a vote of confidence for the reliability of the model. Notably, for marker genes detected in query datasets, the transcriptional profiles of cells assigned to our distinct microglial clusters closely match the profiles of cells in those clusters in our original dataset (Figure S6).

To analyze association of mapped proportion numbers with continuous traits in the ROSMAP single-nucleus data, a linear model from the stats package in R with the formula *Proportion ∼ trait* was used to examine the relationship of amyloid burden to cluster proportion. P-values were adjusted with the Benjamini-Hochberg correction^86^.

### Single nucleus library preparation and sequencing of single nuclei

Dorsolateral Prefrontal Cortex (DLFPC) tissue specimens were received frozen from the Rush Alzheimer’s Disease Center. We observed variability in the morphology of these tissue specimens with differing amounts of gray and white matter and presence of attached meninges. Working on ice throughout, we carefully dissected to remove white matter and meninges, when present. The following steps were also conducted on ice: about 50-100mg of gray matter tissue was transferred into the dounce homogenizer (Sigma Cat No: D8938) with 2mL of NP40 Lysis Buffer [0.1% NP40, 10mM Tris, 146mM NaCl, 1mM CaCl2, 21mM MgCl2, 40U/mL of RNAse inhibitor (Takara: 2313B)]. Tissue was gently dounced while on ice 25 times with Pestle A followed by 25 times with Pestle B, then transferred to a 15mL conical tube. 3mL of PBS + 0.01% BSA (NEB B9000S) and 40U/mL of RNAse inhibitor were added for a final volume of 5mL and then immediately centrifuged with a swing bucket rotor at 500g for 5 mins at 4°C. Samples were processed 2 at a time, the supernatant was removed, and the pellets were set on ice to rest while processing the remaining tissues to complete a batch of 8 samples. The nuclei pellets were then resuspended in 500ml of PBS + 0.01% BSA and 40U/mL of RNAse inhibitor. Nuclei were filtered through 20um pre-separation filters (Miltenyi: 130-101-812) and counted using the Nexcelom Cellometer Vision and a 2.5ug/ul DAPI stain at 1:1 dilution with cellometer cell counting chamber (Nexcelom CHT4-SD100-002). 5,000 nuclei from each of 8 participants were then pooled into one sample, and the 40,000 nuclei in around 15-30ul volume were run on the 10X Single Cell RNA-Seq Platform using the Chromium Single Cell 3’ Reagent Kits version 3. Libraries were made following the manufacturer’s protocol, briefly, single nuclei were partitioned into nanoliter scale Gel Bead-In-EMulsion (GEMs) in the Chromium controller instrument where cDNA share a common 10X barcode from the bead. Amplified cDNA is measured by Qubit HS DNA assay (Thermo Fisher Scientific: Q32851) and quality assessed by BioAnalyzer (Agilent: 5067-4626). This WTA (whole transcriptome amplified) material was diluted to <8ng/ml and processed through v3 library construction, and resulting libraries were quantified again by Qubit and BioAnalzyer. Libraries from 4 channels were pooled and sequenced on 1 lane of Illumina HiSeqX by The Broad Institute’s Genomics Platform, for a target coverage of around 1 million reads per channel.

### Processing of single-nucleus RNAseq reads

For each batch of snRNAseq FASTQ files, the CellRanger software (v6.0.0; 10x Genomics) was used to map reads onto the reference human genome GRCh38, to collapse reads by UMI, and to count UMI per gene per droplet. As a transcriptome model, the “GRCh38-2020-A” file set distributed by 10X Genomics was used. The “--include-introns” option was set to incorporate reads mapped to intronic region of nuclear pre-mRNA into UMI counts. To call cells among the entire droplets, the “remove-background” module of CellBender^95^ was applied to raw UMI count matrices with command line parameters. The admixture of ambient RNA was estimated and subtracted from UMI counts by CellBender. These filtered UMI count matrices were used in the subsequent analyses.

### Demultiplexing

Because our snRNAseq library consisted of nuclei from eight individuals, original individuals of each droplet were inferred by harnessing SNPs in snRNAseq reads. We employed two different procedures, depending on whether all eight individuals had been genotyped with WGS. When eight individuals were genotyped, we used demuxlet^96^ software. From the WGS-based VCF file of 1,196 ROS/MAP individuals, we extracted SNPs that were in transcribed regions, passed a filter of GATK, and at least one of the eight individuals had its alternate allele. The extracted SNP genotype data were fed to demuxlet along with BAM file generated by CellRanger. When less than eight individuals were genotyped, we used freemuxlet (https://github.com/statgen/popscle), which clusters droplets based on SNPs in snRNAseq reads and generates a VCF file of snRNAseq-based genotypes of the clusters. The number of clusters was specified to be eight. The snRNAseq-based VCF file was filtered for genotype quality > 30 and compared with available WGS genotypes using the bcftools gtcheck command. Each WGS-genotyped individual was assigned to one of droplet clusters by visually inspecting a heatmap of the number of discordant SNP sites between snRNAseq and WGS. The above two procedures converged to a table that mapped droplet barcodes onto inferred individuals. Each BAM file generated by CellRanger was split into eight per-individual BAM files, each of which contained reads from distinct individual, using subset-bam (https://github.com/10XGenomics/subset-bam). UMI count matrices filtered by CellBender were split into eight per-individual UMI count matrices.

### Quality control

To identify and exclude potential sample swaps, we assessed concordance of genotypes between snRNAseq and WGS. LOD scores, a metric of genotype concordance, were computed by comparing the per-individual BAM files with WGS genotypes of matched individuals using Picard CrosscheckFingerprints (ver. 2.25.4). We used a haplotype map downloaded from https://github.com/naumanjaved/fingerprint_maps. After inspecting a histogram of LOD scores, ten individuals whose LOD scores were less than 50.0 were filtered out. These individuals received few cells by the demultiplexing procedure. As another measure to detect sample swaps, we checked RNA expression levels of XIST gene and confirmed that they were consistent with clinical sex. Five individuals were further excluded because they failed quality control of WGS. Four were marked as potential sample swaps among WGS, and the other was marked as an outlier of genotype principal component analysis.

Four individual-level sequencing metrics were computed from the per-individual UMI count matrices: estimated number of cells, median UMI counts per cell, median genes per cell, and total genes detected. After inspecting these metrics, individuals whose median UMI counts per cell were less than 1,500 were excluded. Thirteen individuals were found to be sequenced twice in distinct batches. After comparing sequencing metrics, one of these duplicates were excluded from further analyses. After these quality control processes, 424 individuals remained.

### Cell type classifications

To annotate for cell-type, we fitted a weighted ElasticNet-regularized logistic regression classifier over the data of our previous work^97^, predicting for every nucleus one of the 8 major cell types: excitatory neurons, inhibitory neurons, astrocytes, microglia, oligodendrocytes, OPCs, endothelial and pericytes. The gene expression matrix was log-normalized (using NormalizeData method, Seurat package) and scaled over the top 700 variable features excluding non-coding RNA (using FindVariableFeatures method setting the selection method to vst and ScaleData method, Seurat package).

We trained five different models with a mixing parameter of alpha=0 (Ridge), 0.25, 0.5, 0.75 and 1 (Lasso), over a randomly selected 75% of the data (n=139,311). Samples were weighted as 1/ for the number of nuclei of cell-type present in the training set. This step ensured that even low represented cell-types such as endothelial and pericytes will be properly learned. To select the models’ regularization parameters we applied 10-fold cross validation (using cv.glmnet method, glmnet package, 6,7). Fitted models were evaluated using the held out 25% of the data (n=43,428) and their accuracy with respect to the misclassification error was calculated. As all models achieved very high accuracies, with misclassifications mostly between excitatory- and inhibitory neurons, we selected the ElasticNet model with a mixing parameter of \alpha=0.25 to induce sparsity to the model. Fitted model used only 121 out of the 700 available features and achieved a test accuracy of 99.95, with most misclassified nuclei being between inhibitory and excitatory neurons. The nuclei assigned to the microglial cluster were extracted and used in our analyses.

### Leveraging the Connectivity Map to identify chemical and genetic targets for in vitro recapitulation

The Connectivity Map^21, 22^ (CMAP) is a catalog of gene expression signatures for a series of different genetic and pharmacologic perturbations across a wide variety of different cell lines. To identify chemical targets that might drive signatures associated with our distinct microglial subsets in vitro, upregulated gene lists were assembled for each cluster corresponding to genes upregulated in comparison to three or more clusters. The web interface found at clue.io, was used to interface with the CMAP database, and the *ListMaker* tool was used to assemble lists which were then submitted as inputs to the *Query* tool. The version 1.0 L1000 gene expression data compendium was used for all analyses. Output lists were downloaded and ranked by “median_tau_score”. Results were aggregated into families: 1 and 6, 4 and 9, and 8 and 10. Chemical perturbagens of interest were selected from those with a “median_tau_score” above 90 and chosen based on prior knowledge and the pathways they targeted. Full output lists from CMAP separated by cluster can be found in Table S7.

### Drug screening in the HMC3 model system

Compounds of interest were obtained from a wide range of reputable vendors and resuspended in DEPC-treated water (Invitrogen; Cat#:AM9915G), PBS (Corning, Cat #:21-040-CV), or DMSO (Sigma-Aldrich, Cat #:472301). To keep the design of our experiment as similar as possible to the CMAP study, the target stock concentration was 10 mM, but this was adjusted depending on the solubility of each compound. Extensive dose titration with doses ranging from 0.01 µM to 0.1 mM was conducted to determine the highest tolerable dose for each compound. Each concentration of drug was plated in triplicate with early-passage HMC3 cells (ATCC; Cat #: CRL-3304), and the viability was read out using Calcein AM (Invitrogen; Cat #:C1430) and Propidium Iodide (Invitrogen; Cat # : P3566) using a Celigo plate (Nexcelom Bioscience) reader at 6 and 24 hours. An optimal dose of each drug was then chosen based on cell morphology and viability. Subsequently, optimal doses were applied to plated HMC3s and harvested for RNA extraction after for 6 and 24hrs. Lysis was performed in-well with buffer RLT (Qiagen; Cat #: 74136) containing 2-Mercaptoethanol (Thermo Fisher Scientific, Cat #:63689), and RNA extraction was performed with the Qiagen RNEasy mini plus kit (Qiagen; Cat #: 74136) following the manufacturer’s instructions. gDNA eliminator columns were used to remove contaminating genomic DNA. Initial RNA quality and quantity was assessed using Nanodrop (ThermoFisher Scientific) followed by cDNA preparation using the BioRad iScript cDNA Synthesis kit (BioRad; Cat #:1708891). cDNA was subsequently purified with AMPure XP beads (Thermo Fisher Scientific; Cat #: A63880) using a 1:1.8 ratio of cDNA: beads.

### Quantitative RT-PCR analysis

Quantitative real-time PCR reactions to amplify 1 ng of total cDNA were performed in a QuantStudio^TM^ 3 Real-Time PCR Cycler (A28132, Applied Biosystems) using the Applied Biosystems Fast SYBR Green Master Mix (Thermo Fisher Scientific; Cat #: **4385612**). CT values were normalized using Hypoxanthine Phosphoribosyltransferase 1 (*Hprt1*) as the housekeeping gene. Primers were tested for their efficiency beforehand, and the ΔΔC_t_-method was applied for analysis of relative gene expression. The melting curves of each product were analyzed to ensure the specificity of the PCR product.The following primers were used: *HPRT1* -fw: CCTGGCGTCGTGATTAGTGAT, rev: AGACGTTCAGTCCTGTCCATAA; *SRGAP2* - fw: GTTGTGACTTAGGCTACCATGC, rev: TGCTTCGACTGTTCCAGGTTT; *MEF2A* – fw: GGTCTGCCACCTCAGAACTTT, rev: CCCTGGGTTAGTGTAGGACAA; *TYROBP* – fw: ACTGAGACCGAGTCGCCTTAT, rev: ATACGGCCTCTGTGTGTTGAG; *GPX1* – fw: CAGTCGGTGTATGCCTTCTCG, rev: GAGGGACGCCACATTCTCG; *CXCR4* – fw: ACGCCACCAACAGTCAGAG, rev: AGTCGGGAATAGTCAGCAGGA; *SRGN* – fw: GGACTACTCTGGATCAGGCTT, rev: CAAGAGACCTAAGGTTGTCATGG.

### Bulk-RNA Sequencing of compound-treated microglia

0.5×10^6^ HMC3 microglial cells were seeded into a 6-well plate and incubated o.n. The next day, microglia were treated with the respective concentrations of Camptothecin (1µM; EMD Millipore; Cat #:390238), Narciclasine (0.1µM; Millipore Sigma; Cat #: SML2805), Torin2 (10µM; Cayman Chemical Company; Cat #: 14185) or DMSO(Sigma-Aldrich, Cat #:472301) as control and incubated for 24hrs before harvest. Cells were trypsinized (Gen Clone; Cat #:25-510F), counted, the cell viability was assessed and cells were then resuspended in 350µl RLT Lysis buffer (Qiagen; Cat #: 74136) containing 2-Mercaptoethanol (Thermo Fisher Scientific, Cat #:63689), and isolated using Qiagen Plus Mini Kit (Qiagen; Cat #: 74136). RNA quality was assessed using 2100 Bioanalyzer G2938C using an Agilent RNA 6000 Nano Kit (Agilent; Cat # 5067-1511) and Qubit 4 Fluorometer (Invitrogen) using Qubit™ 1X dsDNA HS Assay Kit (Thermo Fisher Scientific, Cat#: Q33231) prior to further processing for RNA sequencing.

mRNA libraries were prepped using Illumina TruSeq Stranded mRNA Library prep (Illumina, Cat#: 20020595), in accordance with manufacturer recommendations, and using IDT for Illumina TruSeq DNA UD Indices (Illumina, Cat#: 20022370) for adapters. Briefly, 500ng of total RNA was used for purification and fragmentation of mRNA. Purified mRNA underwent first and second strand cDNA synthesis. cDNA was then adenylated, ligated to Illumina sequencing adapters, and amplified by PCR (using 10 cycles). The cDNA libraries were quantified using Fragment Analyzer 5300 (Advanced Analytical) kit FA-NGS-HS (Agilent, Cat#: DNF-474-1000) and Spectramax M2 (Molecular Devices) kit Picogreen (Life Technologies, Cat#: P7589). Libraries were sequenced on an Illumina NovaSeq sequencer, using 2 x 100 bp cycles.

Sequencing quality control was performed using Picard version 1.83 and RSeQC version 2.6.1. STAR version 2.5.2a was used to align reads to the GRCh38 genome, using Gencode v25 annotation. Bowtie2 version 2.1.0 was used to measure rRNA abundance. Annotated genes were quantified with featureCounts version 1.4.3-p1.

To analyze the data, a generalized linear model within DESeq2^89^ was used to test for differentially expressed genes across each of our 3 treatment conditions in comparison to control. The DESeq object was constructed with a standard one-factor model, using *∼treatment* as the model for analysis, and genes with less than ten overall counts across all samples were discarded prior to analysis. For analysis of similarity between samples, we used the variance stabilizing transformation in DESeq2, then computed PCA on the resultant matrix. Differential expression was performed with the *DESeq* function, and thresholds for significance were set as an FDR alpha of less than 0.01 and a log_2_fold change (lfc) of 1.5. Shrinkage of lfc was performed with the ashr package^98^, and shrunk lfcs were used for downstream visualization. GO annotation was performed with TopGO^37^ and GO results were summarized with rrvgo^38^. To examine specific genes associated with given cluster families in each treatment condition, the top 20 non-overlapping markers for each member of the grouped clusters (i.e. the top 20 genes for cluster 1, the top 20 genes for cluster 6 that are not in the top 20 gene list for cluster 1) that were present in the differentially expressed gene list for that given condition, regardless of the direction of change (up or down) were chosen for visualization.

### Generation and analysis of global quantitative proteomic data

For global quantitative proteomics of compound-treated HMC3 microglia cells, diaPASEF^99^ (Data independent acquisition) based proteomics was used. In brief, 0.5×10^6^ HMC3 microglial cells were seeded into a 6-well plate and incubated o.n. The next day, microglia were treated with the respective concentrations of Camptothecin (1µM; EMD Millipore; Cat #:**390238**), Narciclasine (0.1µM; Millipore Sigma; Cat #: SML2805), Torin2 (10µM; Cayman Chemical Company; Cat #: **14185**) or DMSO (Sigma-Aldrich, Cat #:472301) as control and incubated for 24hrs before harvest. Cells were trypsinized (Gen Clone; Cat #:25-510F), counted, and the cell viability was assessed. Cells were then washed with ice-cold PBS (Corning, Cat #:21-040-CV) and cellular pellets were snap frozen and store at -80°C until further processing.

Subsequently, cells were lysed in lysis buffer^100^ (1% SDC, 100 mM Tris-HCl pH 8.5, and protease inhibitors; Millipore Sigma, Cat#: D6750; Cat#: 9290-OP) and boiled for 15 min at 60°C, 1500 rpm. Protein reduction and alkylation of cysteine was performed with 10 mM TCEP (Millipore Sigma; Cat#: C4706) and 40 mM CAA (Millipore Sigma; Cat#: C0267)at 45°C for 15 min followed by sonication in a water bath, cooled down to room temperature. Protein digestion was processed for overnight by adding LysC and trypsin in a 1:50 ratio (µg of the enzyme to µg of protein; Promega; Cat#: V5071)) at 37° C and 1400 rpm. Peptides were acidified by adding 1% TFA (ThermoFisher Scientific; Cat#: 28904), vortexed, and subjected to StageTip clean-up via SDB-RPS^100^. Peptides were loaded on one 14-gauge StageTip plug. Peptides were washed two times with 200 µl 1% TFA 99% ethyl acetate (ThermoFisher Scientific; Cat#: 28904; Millipore Sigma; Cat#: 270989) followed 200 µl 0.2% TFA/5%ACN (ThermoFisher Scientific; Cat#: 28904; Thermo Fisher Scientific; Cat#: A955)in a centrifuge at 3000 rpm, followed by elution with 60 µl of 1% Ammonia/50% ACN(Honeywell-Fluka; Cat#: 4427310X1ML; Thermo Fisher Scientific; Cat#: A955) into Eppendorf tubes and dried at 45°C in a SpeedVac centrifuge. Samples were resuspended in 10 μl of LC buffer (3% ACN/0.1% FA) (Fisher Scientific; Cat#: A11710X1-AMP). Peptide concentrations were determined using NanoDrop (ThermoFisher Scientific) and 200 ng of each sample were used for diaPASEF analysis on timsTOFPro (Bruker). Peptides were separated within 120 min at a flow rate of 400 nl/min on a reversed-phase C18 column with an integrated CaptiveSpray Emitter (25 cm x 75µm, 1.6 µm, IonOpticks). Mobile phases A and B were with 0.1% formic acid in water and 0.1% formic acid (Fisher Scientific; Cat#: A11710X1-AMP) in ACN (Thermo Fisher Scientific; Cat#: A955). The fraction of B was linearly increased from 2 to 23% within 90 min, followed by an increase to 35% within 10 min and a further increase to 80% before re-equilibration. The timsTOF Pro (Bruker) was operated in diaPASEF mode^99^ and data was acquired at defined 32 × 25 Th isolation windows from m/z 400 to 1,200. To adapt the MS1 cycle time in diaPASEF, set the repetitions to 2 in the 16-scan diaPASEF scheme. The collision energy was ramped linearly as a function of the mobility from 59 eV at 1/K0=1.6 Vs cm-2 to 20 eV at 1/K0=0.6 Vs cm-2. The acquired diaPASEF raw files were searched with the UniProt Human proteome database in the DIA-NN search engine with default settings of the library-free search algorithm^100^. The false discovery rate (FDR) was set to 1% at the peptide precursor and protein level.

Results obtained from DIA-NN were further analyzed in R. To preliminarily filter the data, peptides without a valid matching gene symbol, as well as peptides that were detected in a fourth of our samples or fewer were removed. For further analyses, total intensity log-normalized protein abundances were used. PCA was performed on the dataset in its entirety to assess relative similarity of treatment conditions. Next, pairwise differential testing between DMSO control and each of our treated conditions was conducted using a Welch’s^92^ t-test with the Benjamini-Hochberg correction^86^, setting a threshold of 0.05 for the corrected p-value and a threshold of 1 for the log_2_ fold change. Top differentially expressed genes were then used for Gene Ontology annotation with topGO (see Bulk-RNA Sequencing of compound-treated microglia). As there were fewer differentially expressed genes overall, all genes associated with each cluster family that overlapped with the differentially expressed gene list for each condition, irrespective of direction (up or down) were selected for plotting.

### Visualizing gene expression across clusters with DotPlots

Seurat’s *DotPlot* function was used to concurrently visualize gene expression and percentage of cells in each cluster expressing said genes. Using this function, a single circle is plotted for each cluster for each given gene. The size of this circle represents the percentage of cells within a cluster that express the gene, and it is absent entirely if fewer than 10% of cells in a given cluster expressed a gene. Conversely, the color of the circle represents the average expression of the gene. This is computed by computing the mean of expression for each cluster, then scaling and zero-centering the average expression level for each discrete cluster. The viridis “magma” color palette was used for this visualization. Legends for the size and color scheme for each dot plot accompany each figure. In addition, for Figure 1C, the “cluster.idents” parameter was used to hierarchically cluster our different clusters by the marker genes involved using complete linkage, enabling clearer visualization of broad differences. The cluster dendogram was manually recomputed and added to the dot plot with the ggtree^103^ package. Notably, data visualization was performed with Seurat v4.0.4 instead of v3.2.0.

### Visualization

All plots were created in R v4.1.0 using either base R visualization packages, ggplot2^104^ with ggrepel^105^, ggfortify^106^, patchwork^107^, cowplot^108^, and ggsci^109^, or packages mentioned in the methods text. Heatmaps were made with the pheatmap^110^ package. Volcano plots were made with the EnhancedVolcano^111^ package.

## Supplemental Information

**Figure S1.**
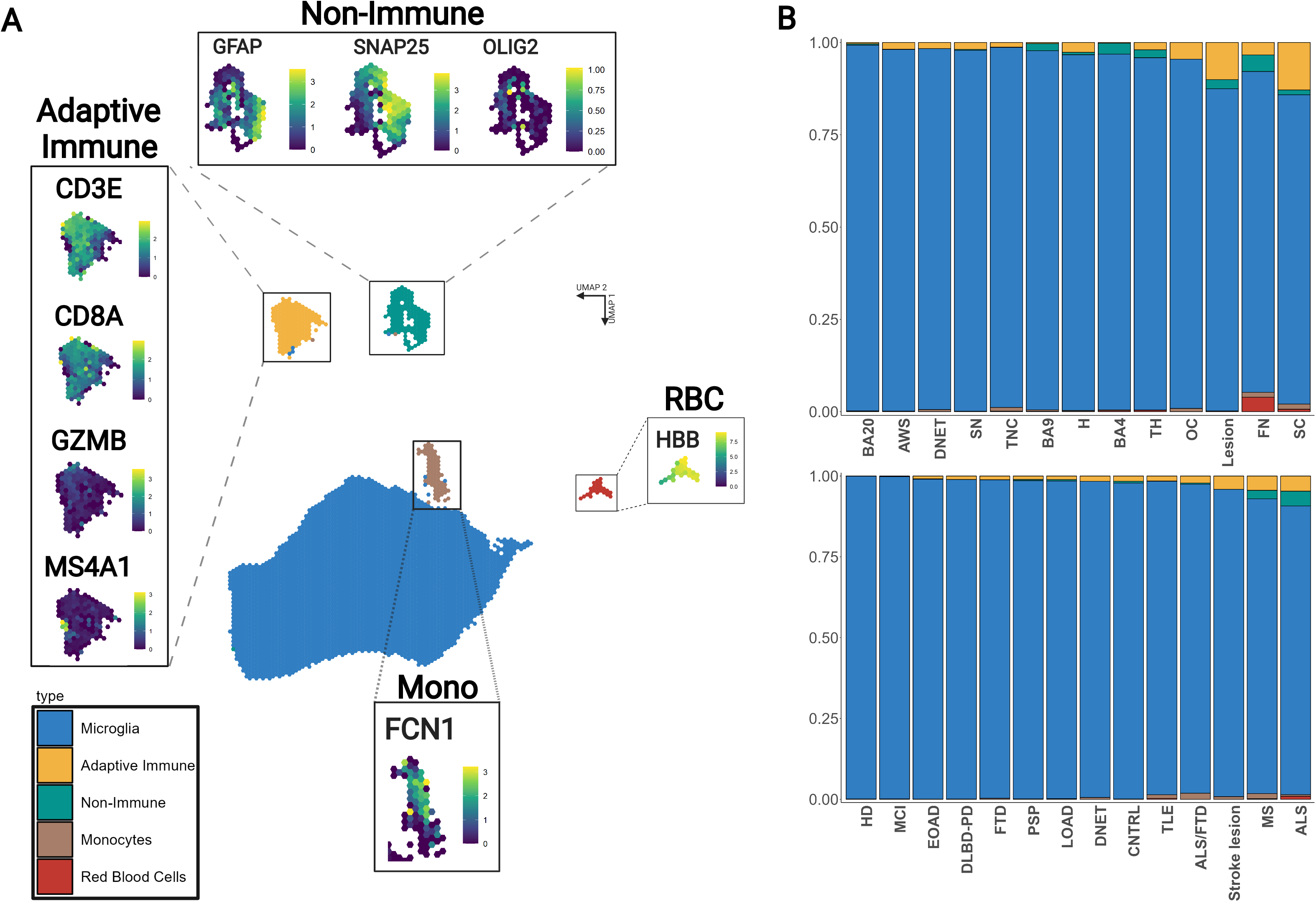
Proportions of overarching cell types in our dataset, related to Methods. **(A) Different cell types are discriminable in UMAP space or by marker genes.** Unsupervised Jaccard-Louvain clustering on a kNN neighbor graph delineates distinct cell types, including adaptive immune cells, monocytes, glial/neuronal cells, and erythrocytes. UMAP plots are binned in hexagons: each single hexagon represents a merged representation of all cells falling within the region. The central UMAP plot is colored by the majority cell type. Different cell types are easily distinguishable in 2-D UMAP plots. The other schex-UMAP plots show gene expression values of selected characteristic marker genes projected onto cells. The color gradient bar represents log-normalized gene expression values. Yellow represents the maximal expressed value, while purple represents the lowest expression values. Markers of distinct immune subpopulations are detected in our data: CD8 T-cells (*CD8A*), NK cells (*GZMB*), B cells (*MS4A1*). Similarly, different non-neuronal cells can be detected in our analysis: astrocytes (*GFAP*), neurons (*SNAP25*), and oligodendrocytes (*OLIG2*). Monocytes (*LYZ*) localize close to our microglial cells and were used for comparative expression of marker genes in Figure 2B. Red blood cells (*HBB)* were also easily discriminable. **(B) Microglia are the predominant cell type recovered across regions and diseases**. Bar plots showing the relative representation of different cell types across different metadata parameters, with each bar summing to 100%. Overall, 95.7% of cells are microglial, 2.2% are adaptive immune, 1.5% are glial/neuronal, 0.4% are monocytic, and 0.3% are erythrocytes. The upper bar plot shows proportion of each overarching cell group across regions, while the lower plot shows the same across diseases. Mono monocytes, RBC red blood cells, LOAD late-onset Alzheimer’s disease, EOAD early onset Alzheimer’s disease, MCI mild cognitive impairment, CNTRL control, DLBD-PD diffuse Lewy body disease-Parkinson’s disease, PSP progressive supranuclear palsy, TLE temporal lobe epilepsy, MS multiple sclerosis, ALS amyotrophic lateral sclerosis, FTD frontotemporal dementia, HD Huntington’s disease, DNET dysembryoplastic neuroepithelial tumor, BA Brodmann area, AWS anterior watershed, OC occipital cortex, TNC temporal neocortex, H hippocampus, TH thalamus, SC spinal cord, SN substantia nigra, FN facial nucleus.

**Figure S2.**
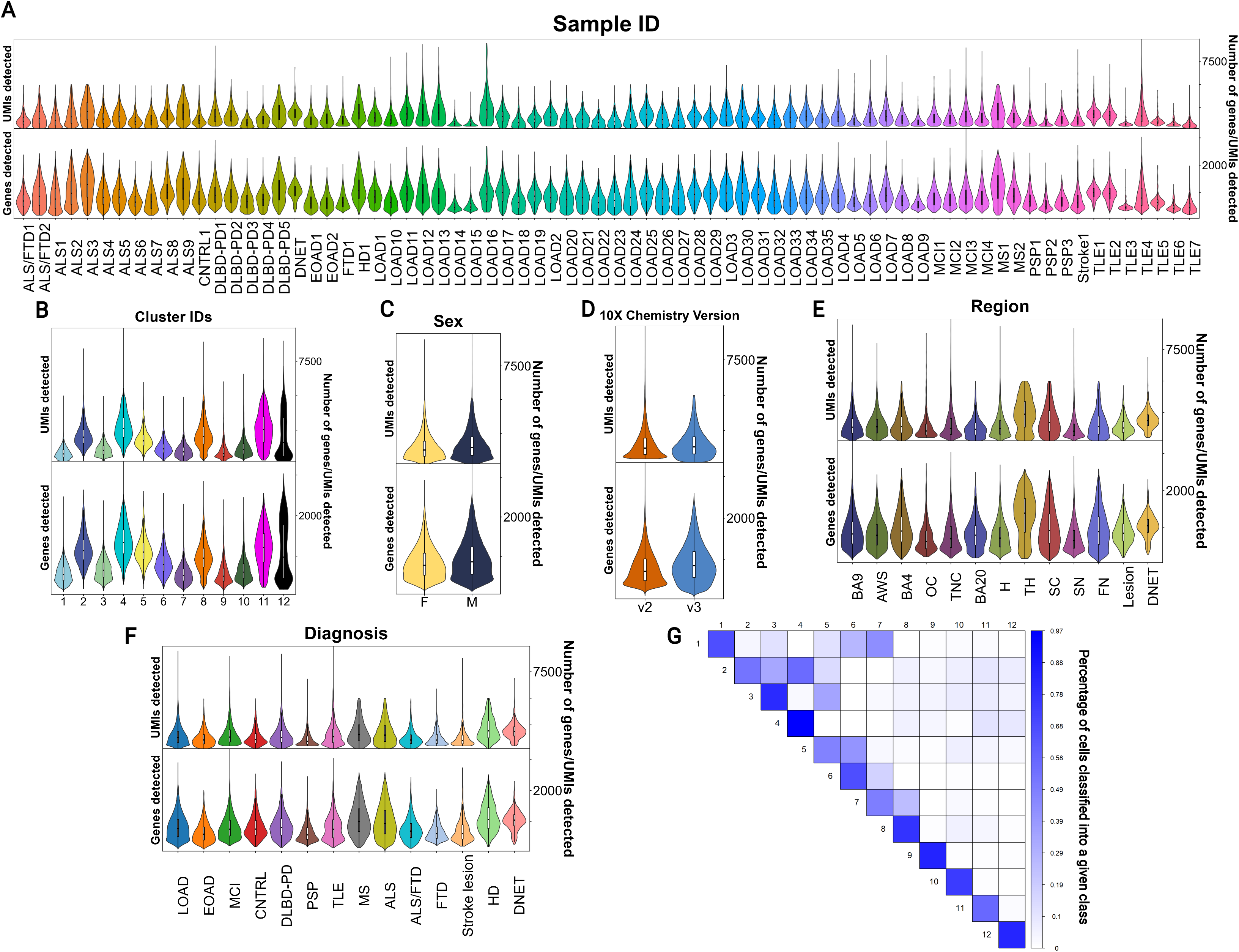
Quality control metrics across our data after downsampling to account for 10x chemistry differences, related to Methods. **(A-D)** Plots are violin plots showing the distribution of the data with overlaid boxplots showing the median, 25% and 75% percentiles. Outliers are not shown in this visualization. Further information about metadata traits may be found in **Supplementary Data 1**. The distributions of unique molecular identifiers (UMIs) and genes detected on a per-cell level after downsampling are similar across donors **(A)**, clusters **(B)**, genders **(C)**, 10x chemistry versions **(D)**, regions, **(E)**, and diagnoses **(F)**. Notably, after downsampling, differences between 10x chemistry versions in these metrics are largely eliminated. **(G) Validation of population stability by resampling and reclustering demonstrates that overlap of gene expression is largely observed for clusters with similarly related families, such as 2 and 4, or for intermediate subsets such as 5 and 3.** To evaluate clustering stability, we randomly sampled ¾ of the cells from our dataset and ran our clustering pipeline with identical parameters. We recorded the frequency of “misclassification”, where cells were re-clustered into clusters different from the one that contained most cells with the same original classification. This process was repeated between pairs of cells, and repeated 50 times for each comparison. Cells were considered to be classified into the “correct” class if they were assigned correctly in ¾ of classification runs. Otherwise, they were considered “misclassified” into a different cluster. Classification frequency is visualized in a heatmap here. LOAD late-onset Alzheimer’s disease, EOAD early onset Alzheimer’s disease, MCI mild cognitive impairment, CNTRL control, DLBD-PD diffuse Lewy body disease-Parkinson’s disease, PSP progressive supranuclear palsy, TLE temporal lobe epilepsy, MS multiple sclerosis, ALS amyotrophic lateral sclerosis, FTD frontotemporal dementia, HD Huntington’s disease, DNET dysembryoplastic neuroepithelial tumor, BA Brodmann area, AWS anterior watershed, OC occipital cortex, TNC temporal neocortex, H hippocampus, TH thalamus, SC spinal cord, SN substantia nigra, FN facial nucleus.

**Figure S3.**
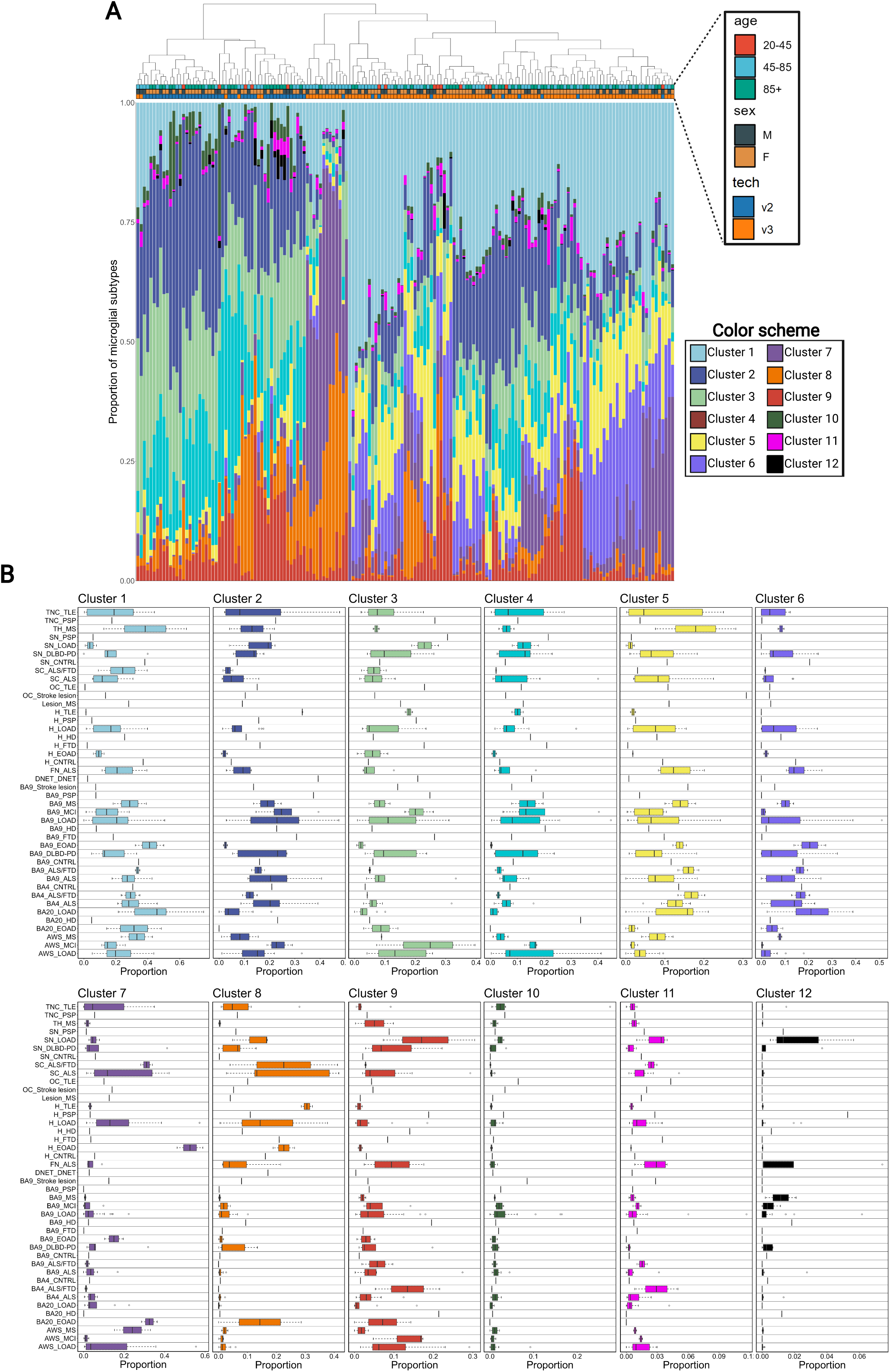
Microglial proportions across individual donors and donor-region pairings. (A) Proportions of microglial subtypes across single donors, related to Figure 3. Proportions of microglial subtypes are plotted by donor, with selected metadata annotated in a header bar above. Each bar represents a single donor and sums to 100%. Samples are clustered hierarchically based on proportions of each subtype. Donors have variability in the exact proportions of different subtypes but exhibit consistent amounts of the most common subtypes in our dataset, clusters 1 through 6. **(B) Proportions of microglial subtypes across region-donor pairings.** Samples are aggregated to donor-region pairings (e.g., AD1-BA9) to give a proportion of different clusters for each region for each individual. Boxplots are computed for specific region-disease pairings showing the median, 25%, and 75% for the proportion of cells across all samples for which that combination of disease and region was sampled. Proportions are shown on the x-axis, and the scale varies depending on the cluster in question.

**Figure S4.**
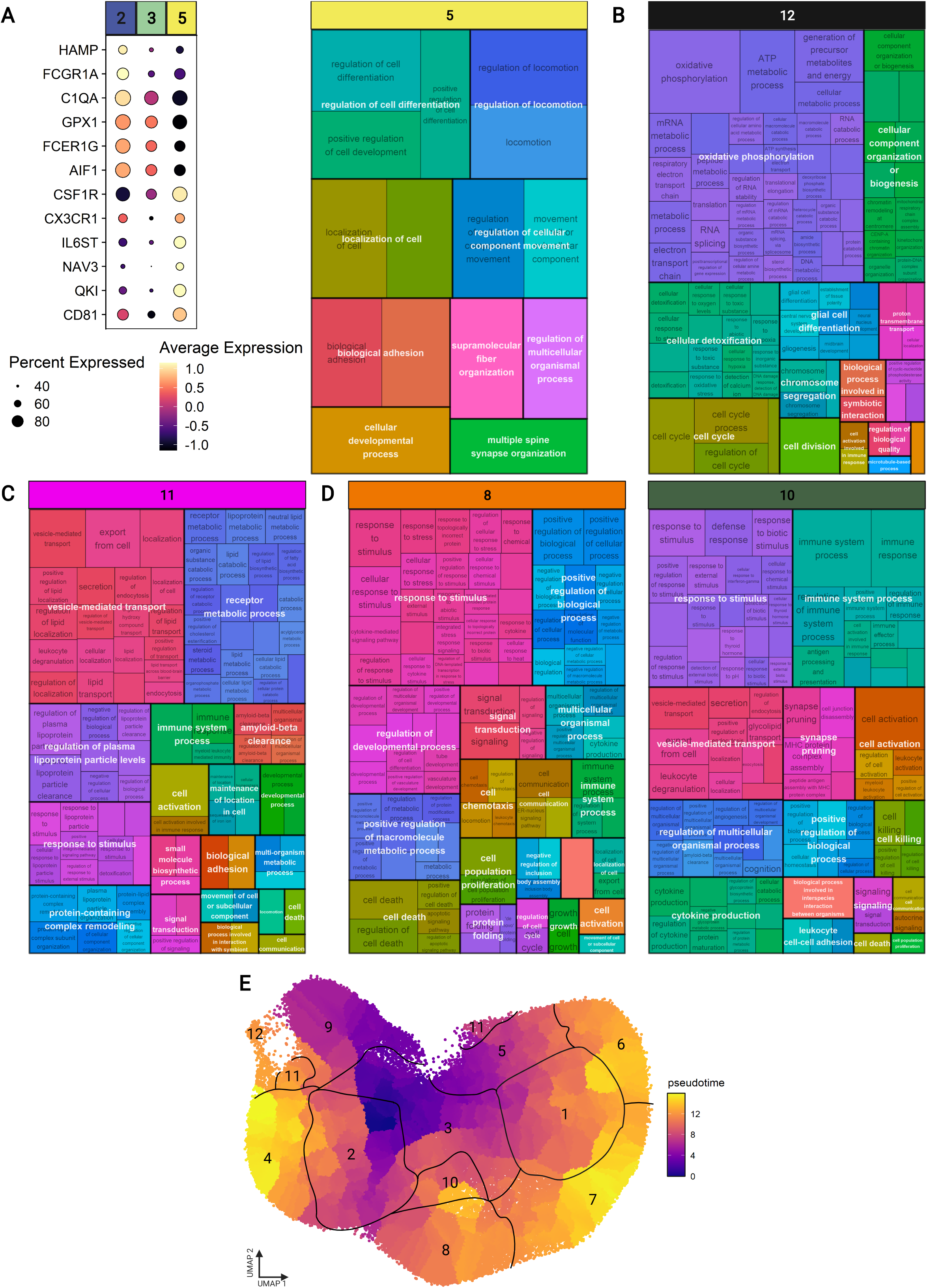
Further exploration of microglial phenotypes with pseudotime analysis and GO annotation validates our trajectory map and reveals subsets associated with motility, lipid trafficking, and proliferation, related to Figure 2. **(A) Cluster 5, an intermediate cluster, shows association with motility.** On the left, the size of the circle represents the percentage of cells in a cluster that express the gene, with no circle plotted if less than 10% of cells in a cluster express the gene. The color of the circle represents the z-scored expression of the gene. Cluster 5 expresses a transcriptional signature partially overlapping with the core homeostatic or transitional clusters, 2 and 3, but expresses unique sets of genes associated with motility. GO annotation was performed with topGO and summarized with rrvgo. Parent terms are shown in white, overlaid over child terms. Terms associated with motility are enriched in cluster 5. **(B) Cluster 12 is associated with oxidative phosphorylation and proliferation. (C) Cluster 11 interfaces with lipids and beta-amyloid. (D) GO annotation of clusters 8/10 parallels results of Reactome pathway analysis, highlighting common immunological activation but divergence in other aspects of phenotype. (E) Trajectories of state shift in pseudotime analysis parallel those seen in other analyses.** Monocle3 was used to build a pseudotime trajectory across our dataset, setting the root point at the boundary of clusters 2 and 3. Shifts in pseudotime from this root point reinforces the directionality laid out in the constellation diagram, suggesting that a broad intermediate gradient between a series of terminal points exists, with pseudotime scores in 6-7, 4, and 10 showing most divergence from the root point. GO gene ontology.

**Figure S5.**
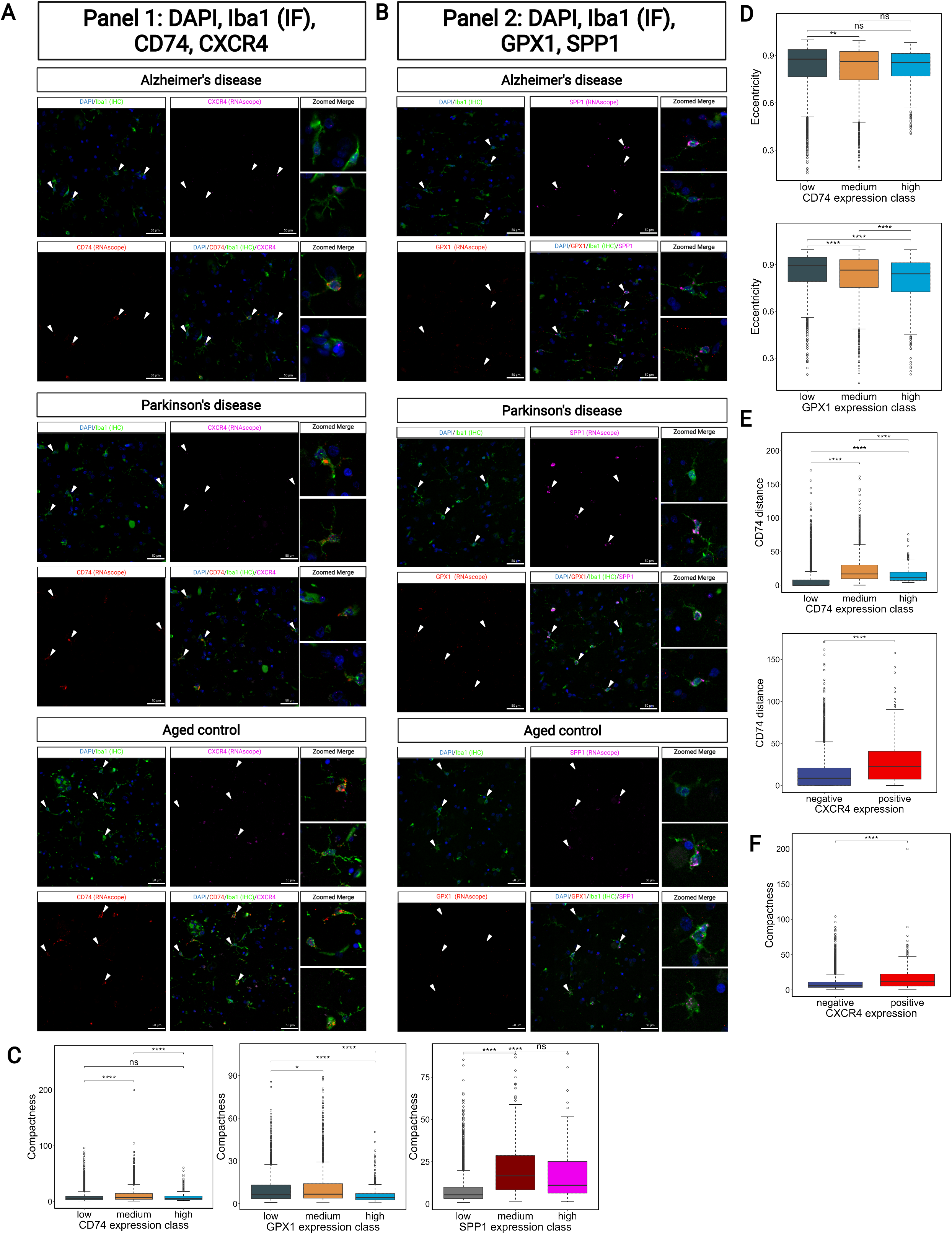
Additional representative images from our joint RNAscope/IF and CellProfiler measures highlight morphological differences between expression-defined subtypes, related to Figure 5. Representative images are shown for both panel 1 **(A)** and panel 2 **(B)** across different diseases. **(C) Compactness is highest in the medium classes of CD74, GPX1, and SPP1-defined expression groups.** Compactness (a measure of ramification, where high values indicate high ramification) is shown across CD74-, GPX1-, and SPP1-expressing Iba1+ microglial cells quantified using CellProfiler. Significance is calculated with two-sample Welch’s t-test. **(D) Compactness is higher in the CXCR4+ class. (E) Eccentricity is highest in the low classes for CD74 and GPX1.** Eccentricity (a measure of shape, where 0 is a circle and 1 is a line), is shown across CD74- and GPX1- expressing Iba1+ microglia. **(F) CD74 distance is highest in the CD74 medium group, but also in the CXCR4^+^ group.** CD74 distance (calculated as the median of all puncta for a given cell from the cellular centroid) is shown across CD74-, and CXCR4-expressing Iba1+ microglia. Significance thresholds: >0.05 = ns, <0.05 = *, <0.01 = **, <0.005 = ***.

**Figure S6.**
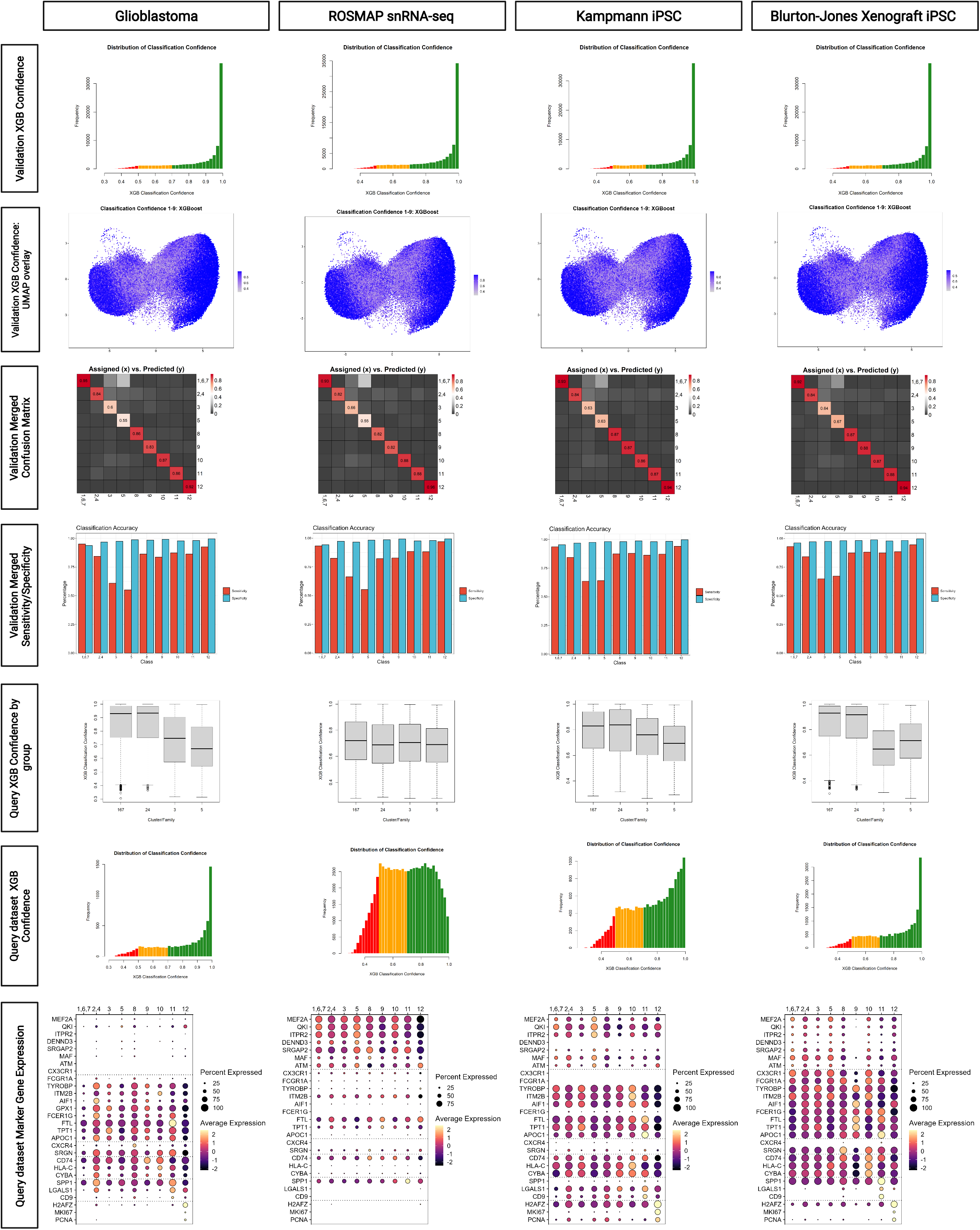
Performance metrics across models trained for different datasets, related to Figure 6. A series of outputs from our label transfer pipeline are presented here. Each row contains a different metric for evaluating performance, while each column represents a single dataset for which a set of models was trained. The training and validation sets for each dataset were identical, but the input differed slightly. For each query dataset we used mNN-corrected data as an input to PCA, but as mNN correction involved the incorporation of the query dataset to correct gene expression, every set of models was trained on slightly divergent input data. The first row presents a histogram of the XGBoost classification confidence output by our models all for cells in the validation set. Cells below 70% confidence are highlighted in yellow, while cells below 50% classification confidence are highlighted in red. Cells below 50% classification confidence were dropped from the model. Most cells in the validation set are classified with high confidence. Row 2 contains a UMAP visualization of classification confidence. It is evident that classification confidence is much higher for cells towards the periphery of the UMAP, whereas intermediate cells in the center of the UMAP have lower overall classification confidence. Row 3 contains the overall confusion matrix for the validation set, while row 4 contains the sensitivity and specificity computed per class. Both are comparable across different query datasets, although they vary slightly. Row 5 contains boxplots for the XGB classification confidence across the 4 classes in question. Boxplots represent the median, 25%, 75% percentiles. Whiskers extend to extremes no more than 1.5 times the interquartile range, and cells that are more extreme than the whiskers are represented as circles. Classification confidence varies substantially depending on the data, with the ROSMAP data being the only dataset where classification confidence for families 167 and 24 was generally comparable to classification confidence for 3 and 5. Row 5 contains histograms of XGBoost classification confidence for all cells in the query dataset. Notably, the glioblastoma and xenograft data have similar classification confidence to the validation set, but the ROSMAP data, and to a lesser extent, the kampmann data, diverge more substantially. Finally, row 6 contains marker gene expression across assigned labels in the query datasets. The size of the circle represents the percentage of cells in a given cluster that express the gene, with no circle plotted if less than 10% of cells in a cluster express the gene. The color of the circle represents the z-scored expression of the gene. Overall levels of expression for these selected genes highlight divergence systematic differences between datasets that persist after batch correction. Nevertheless, label transfer aligns expression profiles effectively.

**Figure S7.**
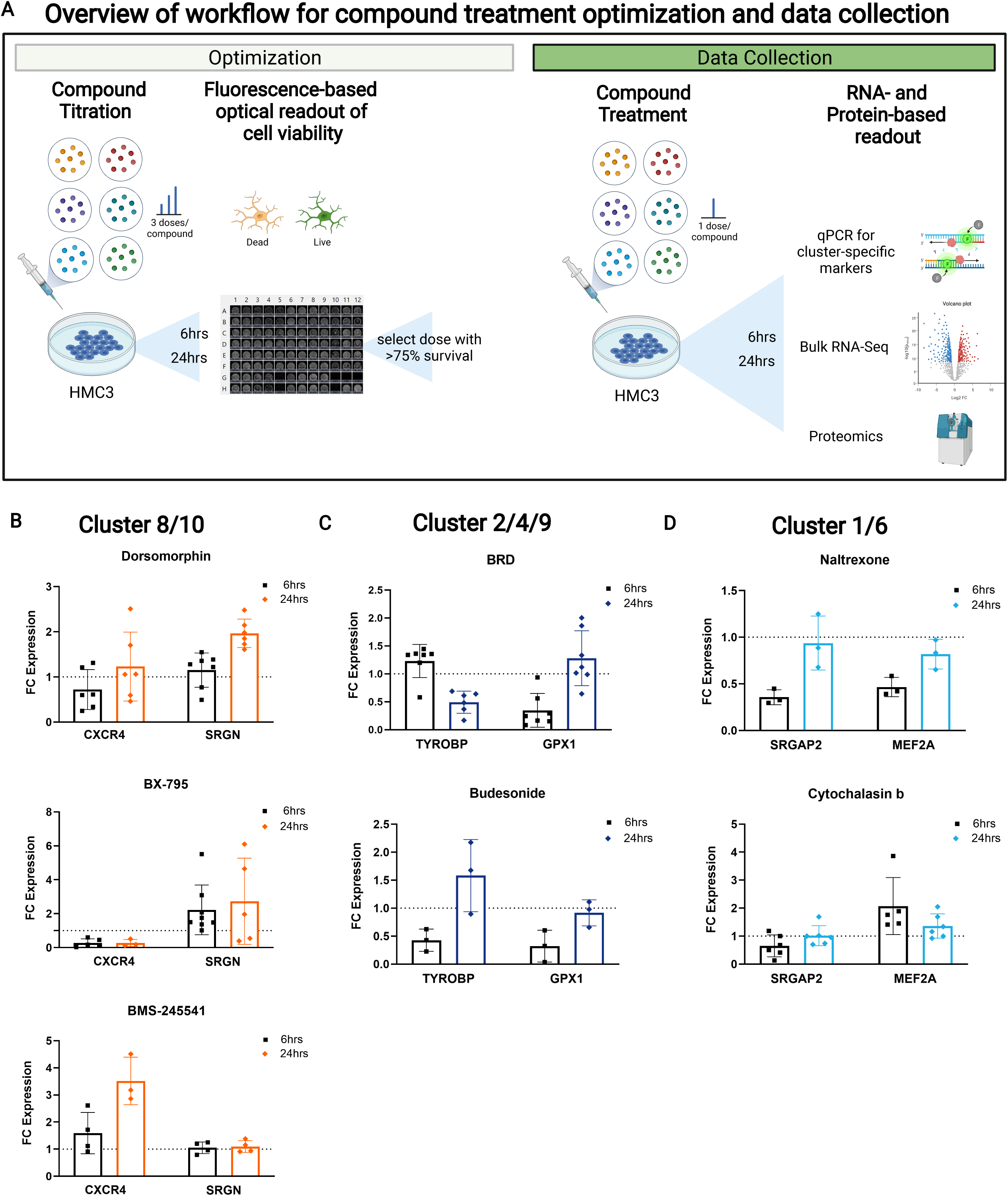
Screening of *in silico* predictions identifies successful hits and compounds that fail to drive predicted signatures. **(A) Schematic overview of workflow for compound treatment, related to Figure 7**. To explore the correct dosage for downstream studies, we conducted dose titration to examine viability of cells after treatment with varying dosages of our drugs. After choosing optimal concentrations, we conducted initial screening with qPCR to select candidates for final validation, then conducted final validation with bulk RNA-seq and proteomics. **(B)-(D) qPCR results for different cluster families.** Results not shown in Figure 10B-D are shown here. Some compounds had effects on specific marker genes, but these did not pass our criteria for further study.

**Figure S8.**
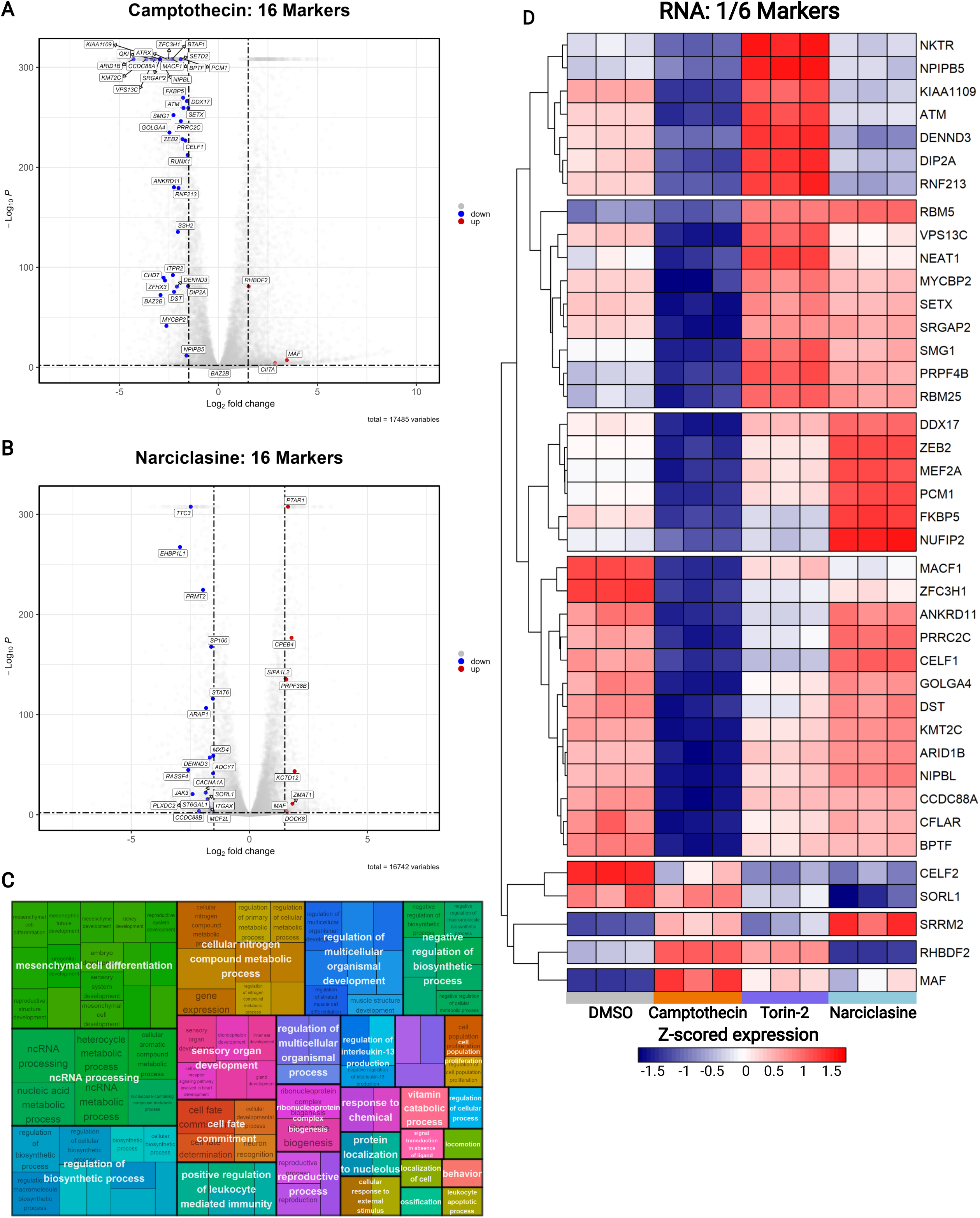
Different compounds modulate different aspects of the cluster 1/6 signature at the transcriptomic level. **(A) Camptothecin downregulates the cluster 1/6 signature, related to Figure 7.** Bulk RNA-seq was generated from HMC3 cells treated with our candidate drugs for 24h. Data was analyzed with DESeq2 and fold change shrinkage was performed with ashr. To examine the genes associated with cluster families, we took the top 20 non-overlapping genes for each individual cluster in our overarching groupings that were present in the differentially expressed gene list for each compound, irrespective of directionality and plotted them in volcano plots. FDR threshold was set to 0.01 and fold change threshold was set at 1.5. **(B) Narciclasine does not upregulate the cluster 1/6 signature. (C) Narciclasine upregulates GO processes also found in cluster 1/6.** GO annotation was computed on differentially expressed genes that passed an FDR threshold of 0.01 and a fold change threshold of 1.5. Terms were grouped based on similar etiology and parent terms were overlaid. Notably, Narciclasine drives metabolic shifts such as in nitrogen-containing metabolism, heterocyclic metabolism, and nucleic acid metabolism, that are strongly enriched in clusters 1/6 (Figure 4A). **(D) Narciclasine and Torin-2 drive distinct modules of cluster 1/6 marker genes.** Cluster 1/6 genes were selected and shown in a row-scaled, zero-centered heatmap. Columns are individual replicates, and rows are genes. These two compounds appear to drive separate modules of genes associated with cluster 1/6. Camptothecin downregulates almost all 1/6 associated genes.

**Table S1 - Overview of demographics, hashing strategy, and quality control, related to Methods.**

**Table S2 - Pairwise marker genes across clusters, related to Figure 1.**

**Table S3 - ROSMAP trait association results, related to Figure 4.**

**Table S4 – Overview of samples used for in situ confirmation, related to Figure 5.**

**Table S5 - CellProfiler pipeline, related to Figure 5.**

**Table S6 - Training results of ML models, related to Figure 6.**

**Table S7 - Results of *in silico* CMAP analysis by cluster, related to Figure 7.**

**Table S8 – *In vitro* validation results at the transcriptomic and proteomic levels, related to Figure 7.**

---

**Graphical Abstract.**
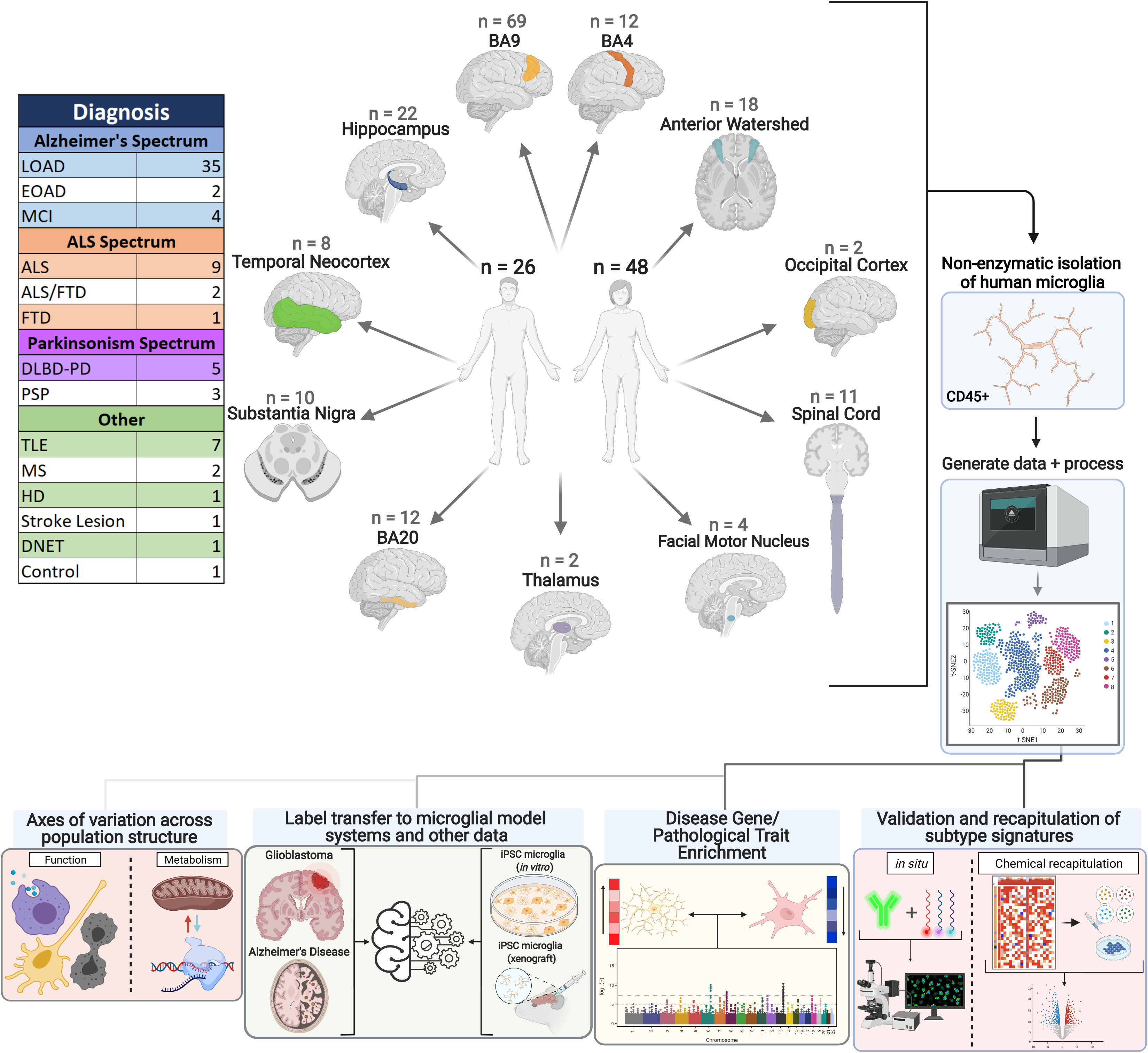
Overview of our cross-disease sample collection, data generation approach, downstream analyses and validation. We sampled a wide array of neurologic diseases and CNS regions (see Table S1) from a mix of autopsy samples and surgical resections. We isolated live brain CD45+ cells from a total of 74 donors of both sexes. Single cell suspensions were loaded directly onto the 10X Chromium controller. Resulting libraries were sequenced on an Illumina Hiseq 4000. The lower part of the figure outlines our analyses and validation efforts, including disease and functional relevance of microglial subtypes, in situ validation, in vitro recapitulation of subtype phenotypes, and annotation of other datasets using our data as a reference.

